# Humanized Extracellular Vesicles for Efficient RNA Delivery

**DOI:** 10.64898/2025.12.15.694436

**Authors:** Xiang Ma, Sophia R. Zhao, Constance L. Cepko

## Abstract

Engineered extracellular vesicles (EVs) are a class of non-viral delivery vectors for RNA-based vaccines and gene therapies. A specialized form of engineered EVs, known as enveloped protein nanocages (EPNs), has been developed to enhance cargo loading and delivery. When EPNs are equipped with a viral fusogen, such as vesicular stomatitis virus glycoprotein (VSV-G), they have been shown to deliver proteins or RNA efficiently into recipient cells. Comparisons across different EPN types and optimization of their different features have been difficult, as assays for their activity have not been reported for single, active units. As we were interested in optimizing EVs, we first developed a biological titration assay inspired by the methods used for infectious viral particles. With this assay, we optimized EVs using a modular platform, creating EVs composed predominantly of human-derived protein components. This system achieved efficient RNA delivery, with functional titers comparable to those of lentiviral vectors. The optimized chimeric proteins comprising the EV particles integrate domains from human epsin 1, human citramalyl-CoA lyase beta-like protein (CLYBL), and human CEP55. The constructs also include a short 21–amino-acid peptide from a non-human source for RNA packaging, resulting in an EV-based RNA delivery system with reduced immunogenicity compared with EPNs and retroviral virus-like particles (VLPs).

**Significance Statement:** We developed engineered extracellular vesicles (EVs) as RNA delivery vehicles to address limitations of virus-like particles (VLPs) and lipid nanoparticles (LNPs) in gene therapies and vaccines. We first developed an assay for individual active particles, using methods typically employed for viral titrations. This approach allowed iterative optimization of a modular EV platform. Our optimized particles comprise primarily human proteins and reach titers that comparable to those of lentiviral vectors.

## Introduction

Delivery of nucleic acids or proteins to recipient cells, particularly in vivo, has historically been done using viral vectors. Viruses evolved many different designs to achieve this, with some of these designs easily adapted for therapeutic or basic science applications. Alternative, non-viral approaches have also been developed. Lipid nanoparticles (LNPs) are a type of non-viral delivery vector for RNA-based vaccines and gene therapies. After entry into cells, ionizable lipids in the LNPs enable release of the RNA payload from the endosome into the cytoplasm. However, only a small fraction of the RNA is successfully released (1–3). It is thus necessary to deliver a high dose of LNPs. The ionizable lipids cause endosomal damage (4) and immune stimulation (5), creating a conflict between efficiency and safety. Extracellular vesicles (EVs) offer an alternative non-viral delivery platform.

EVs are produced by two main pathways: plasma-membrane budding and exocytosis of multivesicular bodies, whose fusion with the plasma membrane releases exosomes (6). RNA can be loaded into EVs endogenously during their biogenesis in producer cells. Alternatively, RNA can be loaded into isolated EVs exogenously by transiently permeabilizing EV membranes with physical or chemical methods (7). Even without a known fusogen, RNA-loaded EVs can be internalized by recipient cells and elicit functional responses; however, delivery is typically inefficient, and the underlying mechanisms remain incompletely characterized (8, 9).

Engineered EVs and other engineered particles, such as retrovirus-based VLPs, can incorporate well-characterized fusogenic viral glycoproteins, such as VSV-G. Such glycoproteins drive endocytosis and low pH-triggered membrane fusion within endosomes, by pulling together the viral and endosomal membranes (10). This mechanism is distinct from the membrane rupture associated with ionizable lipids. It is often used by enveloped viruses to liberate viral components into the cytoplasm and, in engineered particles, to promote payload delivery (11–15). Enveloped protein nanocages (EPNs) are a type of engineered EV that have been shown to package and transfer proteins or RNA between cells (11, 16). EPNs carrying Cre mRNA have been assayed by quantifying the percentage of reporter-positive recipient cells using flow cytometry (11). However, such studies did not report biological titers, as one typically does to quantify active viral particles, limiting comparisons among constructs, batches, and delivery systems.

We wished to engineer efficient and non-toxic EVs for RNA delivery in therapeutic applications. To accomplish this, we first developed a biological activity assay inspired by methods used to measure active viral particles. We used this method in combination with measurements of EV-packaged RNA, as well as translation of mRNA cargo in recipient cells. After multiple rounds of optimization, we established a modular EV platform for RNA delivery composed predominantly of human-derived proteins, achieving functional titers comparable to those of lentiviral vectors (17). As with the pioneering designs of Votteler et al. and Horns et al. for EPNs, we created EVs from a single chimeric protein integrating programmable modules (11, 16). Our designs use: (i) a membrane-targeting domain—either the rat phospholipase C delta 1 pleckstrin homology (PLCδ1-PH) domain or the human epsin-1 epsin N-terminal homology (ENTH) domain; (ii) a multimeric scaffold—the bacterial 2-keto-3-deoxy-6-phosphogluconate (KDPG) aldolase, the I3-01 protein nanocage, or the human citramalyl-CoA lyase beta-like protein (CLYBL) trimer; (iii) an ESCRT (endosomal sorting complex required for transport) recruitment domain using human CEP55 ESCRT/ALIX-binding region (EABR) or HIV-1 Gag p6 peptide, and (iv) an RNA tethering domain using lambda bacteriophage N (λN) peptide. Initially, we incorporated a drug-controlled HCV NS3/4A protease to cleave the λN-tethered RNA payload from the multimeric scaffold of the particle, after entry into recipient cells, thereby retaining high-avidity loading in producer cells while minimizing translational drag in recipients. By examining the contributions of each of these modules, we found that nanocage formation was not essential for efficient production and delivery of functional EVs, whereas λN valency was a dominant determinant. To reduce immunogenicity, we humanized the scaffold (substituting CLYBL for the bacterial nanocage) and removed the HCV NS3/4A protease; to sharpen plasma-membrane targeting, we replaced rat PLCδ1-PH with human epsin-1 ENTH (S4W). Ultimately, these optimizations enabled the design of an efficient EV particle along with a paired cargo plasmid encoding an RNA payload that is robustly packaged into EVs and efficiently translated in recipient cells.

## Results

### Modular design of EVs for RNA delivery

We first developed a biological titration assay to quantify functional delivery units per milliliter (DU/mL). This assay is based on titration methods for lentiviral vectors and recombinant VSVs (18, 19). Baby hamster kidney (BHK-21) cells served as the recipient cells for EVs produced by 293T producer cells. BHK-21 cells were transiently transfected with a Cre-dependent DsRed reporter (pCALSL-DsRed). Serial 10-fold dilutions of EV-containing supernatant from 293T producer cells were then added to a series of BHK-21 culture wells, which were cultured for up to 4 days (Fig. 1A). Compared with a Cre-reporter cell line containing genomically integrated transgenes for detecting Cre mRNA delivery (11, 12), transiently transfected recipients contained multiple plasmid copies. This resulted in bright, readily counted DsRed-positive cells after Cre–lox recombination, enabling direct quantification with an inverted fluorescence microscope.

**Fig. 1.**
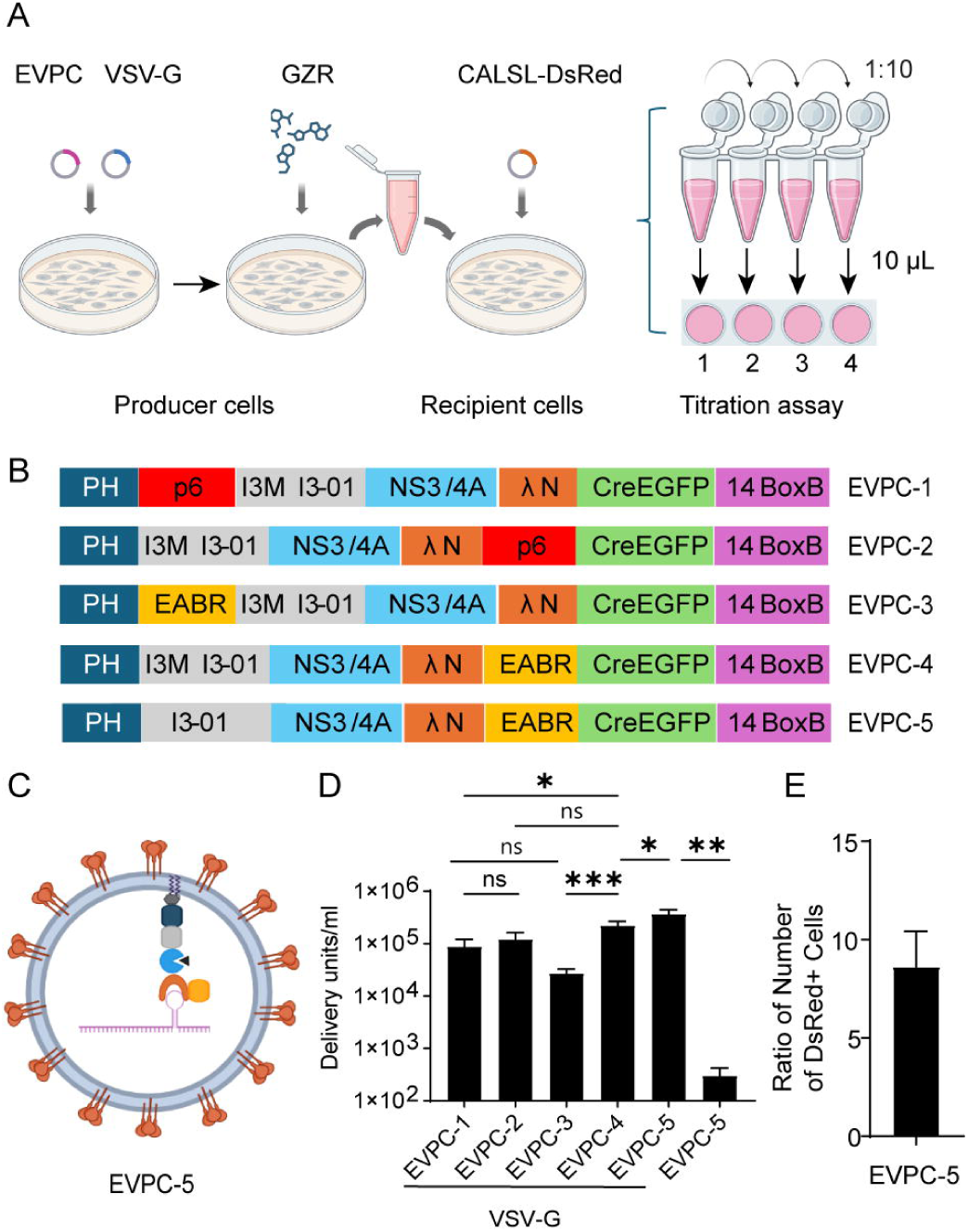
Strategy of EV production and their titration. (A) Biological titration assay workflow. 293T producer cells are transfected with EV packaging-cargo construct (EVPCs) and VSV-G, then cultured in the presence of the HCV NS3/4A inhibitor grazoprevir (GZR). Cell-free supernatant is serially diluted 10-fold; 10 µL of each dilution is added to BHK-21 recipient cells (transiently transfected with pCALSL-DsRed) in 500 µL of medium (first well, 10 µL undiluted supernatant). After 4 days, DsRed⁺ cells are counted by inverted fluorescence microscopy. (B) EVPC design. All variants contain the PLCδ1-PH membrane-targeting domain, the I3-01 self-assembling nanocage (with Met3 in EVPC-1 to EVPC-4; reverted to Ile in EVPC-5), HCV NS3/4A protease, and a λN peptide (ΔMet1) for BoxB-mediated mRNA tethering; Cre-P2A-EGFP is linked downstream via T2A, and a 14×BoxB array is placed in the 3′UTR. ESCRT-recruitment modules are either HIV-1 Gag p6 or the CEP55 EABR, positioned internally (between PH and I3-01) or at the C terminus. (C, D) Schematic of EVs produced by coexpression of EVPC-5 and VSV-G plasmids, with the functional titers plotted (D) as DU/mL (mean ± SEM, n = 8). *p*-values across EVPC-1 to EVPC-4 were calculated using Dunnett’s multiple comparisons test following the Friedman test. *p*-values between EVPC-4 and EPVC-5 were calculated using paired t-test. *p*-values for comparisons with or without VSV-G, in combination with EVPC-5, were calculated using the Wilcoxon test. (E) Test of linearity of titration method: adjacent wells had serial 10-fold dilutions of an EV supernatant added, with counts of DsRed⁺ cells carried out 4 days later. The fold change among wells was measured (n = 8) and the *p*-value was calculated using a one-sample t-test against a value of 10.

To optimize the EV design for RNA delivery, we compared biological titers from constructs varying three elements of EV production: plasma membrane-targeting domains, self-assembly components, and modules that recruit the ESCRT pathway to promote EV release (16). Horns et al. designed EPN24-MCP, which had the nanocage component I3-01 fused to the RNA-binding domain derived from MS2 bacteriophage coat protein (MCP), creating a self-assembling nanocage that binds RNA with the cognate MS2 hairpin (11). We designed four variants of EPN24-MCP, named EVPC-1 to EVPC-4, by constructing EV packaging–cargo (EVPC) plasmids that combine the EV packaging and RNA cargo functions within single vector (Fig. 1B). All constructs retained the plasma membrane-binding PH domain of PLCδ1 and the self-assembling nanocage I3-01. A previous study showed that a methionine codon at position 3 of I3-01 could serve as an internal translation initiation site (16). We hypothesized that nanocage protomers translated from this internal start codon and thus bearing a free N-terminus, might facilitate nanocage assembly and enhance EV production. To test this, we mutated the isoleucine codon at position 3 of I3-01 to methionine.

For ESCRT-recruiting components, both the HIV-1 Gag p6 peptide (16) and EABR fragment of human CEP55 were tested. The EABR fragment has higher binding affinity to ALIX and the ESCRT-I subunit TSG101 than viral late domains such as the HIV-1 p6 (20). This fragment has been shown to facilitate greater release of enveloped VLPs (eVLPs) when fused to the SARS-CoV-2 spike cytoplasmic tail (21). We hypothesized that constructs with EABR could generate higher titers of EVs than constructs with the p6 peptide. Although the p6 peptide is naturally located at the C terminus of the HIV Gag protein, it remains functional when positioned internally (22). In contrast, the EABR fragment, which is naturally located internally within CEP55 (20), remained functional when placed at the C terminus of membrane proteins (21, 23). To investigate the positional dependence of these elements, we designed four constructs in which either the p6 peptide or the EABR domain was placed at the C terminus or inserted internally between the PH domain and the I3-01 nanocage (Fig. 1B).

To load cargo mRNA into EVs in producer cells, the MCP used in a previous study by Horns et al. (11) was replaced by the λN peptide. The short 22-amino-acid λN peptide has been used to tag proteins (e.g., GFP). When its specific 19-nucleotide binding site (BoxB) is inserted into a target RNA, the high-affinity interaction between λN and BoxB enables tethering of proteins to RNA or vice versa (24–30). The main advantage of the λN peptide relative to MCP is its small size and monomeric nature (31, 32), which reduces potential interference between the EV protein chimera and the tethered RNA. Moreover, multiple copies of λN could be fused in tandem within the EV protein chimera to enhance its binding avidity to the target RNA (26, 29). The first methionine is dispensable for BoxB binding (32–34); therefore, the initial methionine of the λN peptide was deleted to prevent internal translation initiation. As a result, only 21 amino acids of this peptide were included in the constructs.

To deliver cargo RNA into recipient cells, VSV-G was used as a fusogen as in previous studies (11, 16). To enhance endosomal escape and potentially increase RNA translation efficiency in recipient cells, a drug-controlled hepatitis C virus (HCV) NS3/4A protease together with its cognate cleavage site was included in the construct (35–39). This protease was designed to cleave between the λN peptide and the PH domain-nanocage inside recipient cells, with the hypothesis that the liberated RNA would be more efficiently translated. In producer cells, a small-molecule drug was used to inhibit protease activity, allowing efficient RNA loading and EV production. Upon transfer to recipient cells, the inhibitor was diluted, enabling the protease to cleave at its C terminus and release the λN-RNA payload (SI Appendix, Fig. S1A).

The initial constructs encoded a large Cre-containing mRNA cargo (∼5□kb), in which Cre-P2A-EGFP was linked to EPN24-λN variants via a T2A sequence, with a 14x BoxB array positioned in the 3’ UTR. P2A (porcine teschovirus-1 2A) and T2A (Thosea asigna virus 2A) peptide linkers mediated ribosomal skipping (40), enabling coexpression of all three components (Fig. 1B). In producer cells, EGFP fluorescence served as an indicator of transfection efficiency. When coexpressed with VSV-G in 293T cells in the presence of HCV NS3/4A protease inhibitor, the EPN24-λN variants packaged their own mRNA, which was then delivered by the EVs into BHK-21 cells. EVPC-4 yielded the highest titer, significantly exceeding EVPC-1 and EVPC-3 but not EVPC-2. Constructs containing the p6 peptide (EVPC-1, −2) produced similar titers, whereas those with the EABR domain (EVPC-3, −4) differed, with EVPC-3 generating significantly fewer EVs. Despite the higher affinity of EABR for ESCRT components, substituting it for p6 in either position did not significantly alter EV output. In pilot tests, placing EABR between I3-01 and NS3/4A reduced titers and was not pursued. To assess whether an isoleucine-to-methionine substitution at position 3 of I3-01 influenced EV production, we reverted the methionine to isoleucine in EVPC-4. The resulting construct, EVPC-5, showed a statistically significant increase in EV production efficiency. When EVPC-5 was transfected into 293T cells without cotransfection of VSV-G plasmid, almost no DsRed+ cells were detected among the recipient cells (Fig. 1 C and D; SI Appendix, Fig. S1B). However, RT-qPCR analysis of the supernatant from producer cells showed that Cre mRNA export was not diminished (SI Appendix, Fig. S1C). These results confirm that successful mRNA delivery requires VSV-G-mediated endosomal escape of EVs.

Based on previous studies (38), the HCV NS3/4A protease inhibitor Grazoprevir (GZR) was tested at 1,250□nM, along with additional concentrations, to evaluate how GZR levels in producer cells affected the EV titers generated by EVPC-5. Increasing the GZR concentration two- or fourfold allowed an assessment of the concentration required to inhibit NS3/4A protease activity and enable mRNA packaging and EV release. At the same time, we considered the need for GZR to be sufficiently diluted in recipient cells to permit protease-mediated cleavage and release of the λN-mRNA payload. In the titration assay, recipient cells were exposed to serial 10-fold dilutions of EV-containing supernatant to quantify functional delivery. Because the first well in this dilution series contained 10□μL of supernatant added to 500□μL of culture medium, this is a 50-fold dilution of GZR. To assess whether residual GZR inhibited NS3/4A activity in recipient cells, 10- and 50-fold reductions in GZR concentration were tested to determine their effects on EV titers. When no GZR was added to producer cells, EVs were released at 1,800 DU/mL. We hypothesized that these EVs reflected passive mRNA packaging and release, independent of λN-BoxB binding and EABR function. In contrast, addition of 1,250□nM GZR enabled active mRNA loading and ESCRT-dependent EV release, resulting in a marked increase in titer. Higher GZR concentration did not further enhance EV titers. Reducing the GZR concentration to 125 nM significantly decreased EV titers, whereas 25 nM GZR resulted in titers comparable to the no-GZR condition (Fig. 1A; SI Appendix, Fig. S1D). These results suggested that 1,250□nM GZR is sufficient to inhibit protease activity in producer cells but was diluted in the first well of the titration assay, permitting protease-mediated cleavage. Consequently, subsequent wells in the 10-fold dilution series also fell below the inhibitory threshold.

In the titration assay, we counted DsRed+ cells in wells containing fewer than 2,000 positive cells, corresponding to a multiplicity of infection (MOI) of less than 0.01. We quantified DsRed+ cells in neighboring wells, which received 10-fold dilutions of EV-containing supernatant produced by cotransfection of EVPC-5 and the VSV-G plasmid. The ratio of DsRed⁺ cell counts between adjacent wells was 8.6-fold, not significantly different from the theoretical value of 10 (*p*□= 0.48) (Fig. 1E). This linear decrease in delivery units across dilutions supports the assumption that each DsRed⁺ cell arose from a single functional EV-mediated delivery event of Cre mRNA, allowing accurate back-calculation of functional EV titers.

### Loss-of-function variants reveal domain requirements for functional EV production

We next generated multiple deletion and point mutation variants to determine which components of the EVPC-5 construct were essential for functional EV production. The EVPC-5 +VSV-G group, comprising 45 batches, had a titer 3.5 × 10□ DU/mL and served as the positive control. The EVPC-5 construct, expressed without cotransfection with the VSV-G plasmid, yielded a titer of 260 DU/mL across 37 batches and served as negative control (Fig. 2A).

**Fig. 2.**
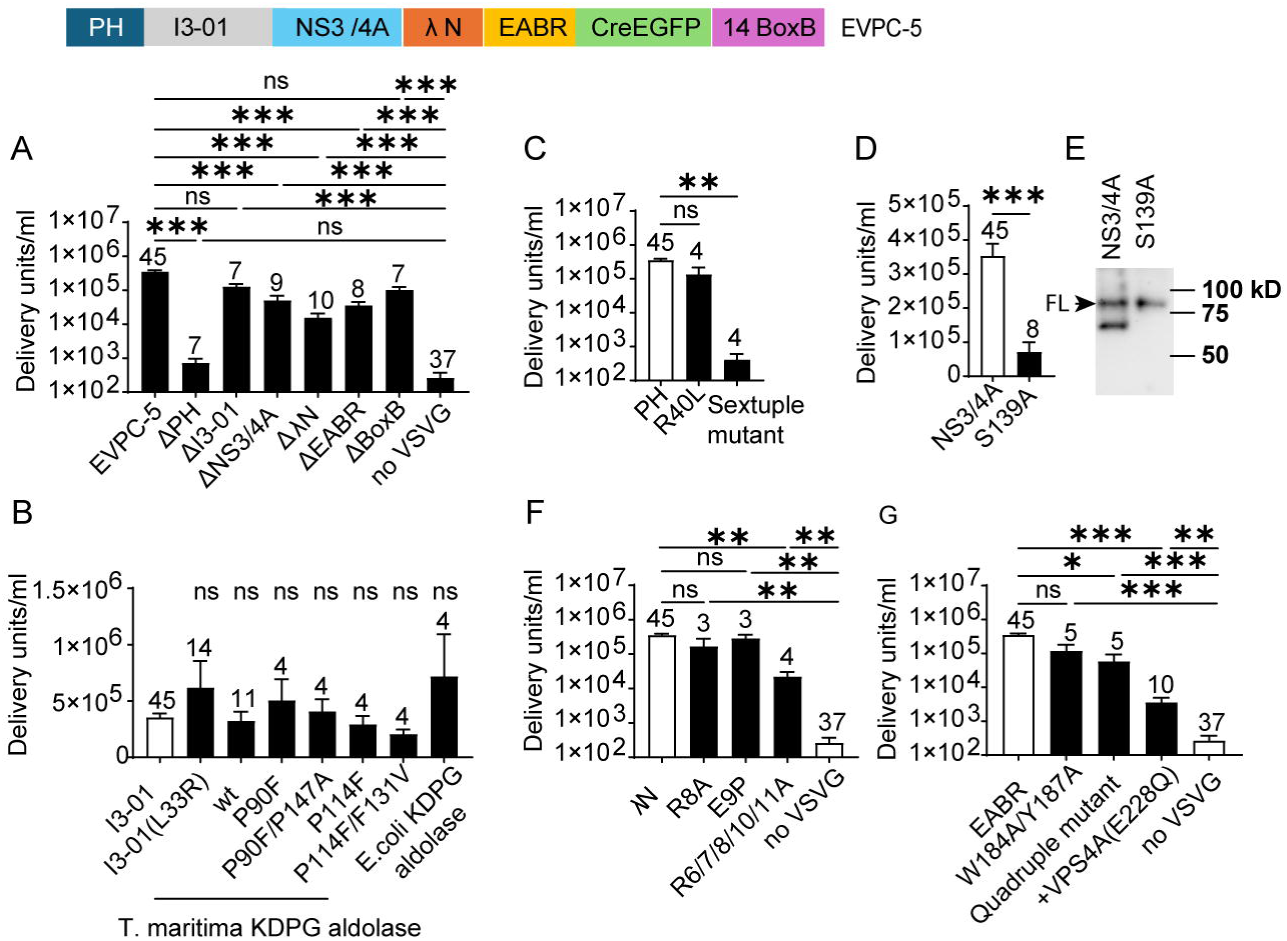
Modular requirements for functional EV production in the EVPC-5 system. (A) Domain dependence assessed by single-component deletions, including positive control (EVPC-5 + VSV-G), non-VSV-G control (EVPC-5 without VSV-G), and six single-component deletion variants. (B) Oligomerization requirement assessed by replacing the I3-01 nanocage with I3-01 protomer L33R point mutant (trimer), wild-type trimeric KDPG aldolase (*T. maritima* or *E. coli*) or by using *T. maritima* KDPG aldolase monomeric variants (P90F, P114F) and their corresponding variants reverted to the trimer form (P90F/P147A, P114F/F131V). (C) Test of requirement for PLCδ-PH-mediated plasma membrane targeting assessed by R40L and a six-basic-residue (Sextuple) mutant. (D) Test of requirement of NS3/4A protease activity assessed by a protease-deficient mutant (S139A). (E) Test of cleavage of EV chimeric proteins by NS3/4A protease. EV-containing supernatants were centrifuged, then incubated in inhibitor-free medium to permit limited NS3/4A cleavage. Cleavage was detected by western blot using an anti-Myc antibody (tag on I3-01). Uncleaved and cleaved protein bands of the EV chimeric proteins are shown. Representative blot, n = 3. (F) Requirement for λN was assessed with R8A, E9P, and five-arginine-to-alanine (R6/7/8/10/11A) substitutions within the λN RNA-binding domain. (G) Test of requirement for ESCRT interactions via EABR. Dependence was assessed using W184A/Y187A and W184A/Y187A/R191A/E192A (Quadruple) substitutions within EABR, as well as by coexpressing dominant-negative VPS4A (E228Q) with EVPC-5 and VSV-G. Nonparametric tests were applied when comparing either the positive or non-VSV-G control groups with EVPC-5 variants. *p*-values were calculated using the Kruskal–Wallis test followed by Dunn’s multiple comparisons test, except in (D), where a Mann–Whitney test was used.

#### Deletion or loss-of-function variants of membrane-targeting PLCδ1-PH domain

Deletion of the plasma membrane-targeting PLCδ1-PH domain reduced EV production to 730 DU/mL, a 485-fold decrease to a level comparable to the non-VSV-G control (Fig. 2A). Surprisingly, the R40L point mutation, which disrupts ligand binding and plasma membrane localization of PLCδ1-PH-GFP (41, 42), did not abolish EV production. However, substitution of six basic residues (K30L, K32L, R37D, R38E, R40A, and K57A) in the PLCδ1-PH domain that are critical for ligand binding (43) reduced EV titers to 400 DU/mL, comparable to the ΔPH variant (Fig. 2C). We hypothesized that positioning the PH domain at the N-terminus of oligomerization scaffold I3-01 enhances its avidity for PI(4,5)P₂, such that more extensive disruption of critical basic residues is required to fully prevent membrane targeting of PH-I3-01 than in PH-GFP.

#### Deletion or loss-of-function variants of self-assembly component I3-01

Deletion of the self-assembly component I3-01 reduced EV titers by 2.7-fold, which was not statistically significant compared with the positive control (*p* = 0.33) (Fig. 2A). Likewise, the L33R point mutation, which disrupts nanocage assembly and has been reported to abolish EV production in the EPN-01 design (16, 44), produced EVs with titers comparable to the positive control. The I3-01 nanocage comprises 60 subunits and is derived from trimeric KDPG aldolase from *Thermotoga maritima (*PDB ID: 1WA3). The protomer contains five substitutions (E26K, E33L, K61M, D187V, and R190A) relative to the wild-type enzyme. Notably, the L33R mutation reverts the I3-01 nanocage to a trimeric form (44, 45). To further test whether nanocage assembly is required for EV production, we replaced I3-01 with the wild-type *T. maritima* KDPG aldolase, which normally forms a trimer. This construct produced EVs at titers not significantly different from I3-01 or its L33R mutant. Similarly, replacing I3-01 with the trimeric *E. coli* KDPG aldolase (46) yielded comparable titers. The wild-type *T. maritima* KDPG aldolase can be converted to a monomer by single-residue substitutions, and compensatory mutations restore its trimeric assembly (47). To further probe the role of the oligomeric state, we tested monomeric variants (P90F, P114F) and reverted trimeric variants (P90F/P147A, P114F/F131V), all of which produced EV titers comparable to I3-01 (Fig. 2B). These results demonstrate that within EVPC-5, the 60 subunit-nanocage is not essential for functional EV production; trimeric and even monomeric variants of *E.coli* and *T. maritima* KDPG aldolase support EV production at levels equivalent to the original I3-01 construct.

#### Deletion or loss-of-function variants of the NS3/4A protease

Deletion of the NS3/4A protease reduced EV production by 7-fold but remained significantly higher than the non-VSV-G control (Fig. 2A). The HCV NS3 protease is a serine protease that requires the NS4A peptide as a cofactor. Serine 139 is a critical residue in its catalytic triad; substitution with alanine (S139A) abolishes protease activity (48–51). A construct carrying the S139A mutation produced EVs at 5-fold lower titer compared with the wild-type NS3/4A construct (Fig. 2D). To verify protease activity within EVPC-5, EV-containing supernatants were concentrated by ultracentrifugation and incubated at 37 °C for 30 min in fresh medium lacking GZR to permit protease-mediated intra-EV cleavage. Because of the brief incubation following inhibitor removal, only partial cleavage was expected. Notably, in the titration assay, recipient cells were analyzed 4 days after EV addition, providing additional time for intracellular cleavage. The S139A mutant served as an uncleaved control. Western blotting with an anti-Myc antibody (targeting the Myc-tagged I3-01 domain) detected both full-length and cleaved protein in the wild-type EVPC-5 group, whereas only the full-length band was observed in the S139A mutant (Fig. 2E).

#### Deletion or loss-of-function variants of the λN domain

Deletion of the λN domain reduced EV titers 23-fold but remained significantly higher than the non-VSV-G control (Fig. 2A). Single-point mutations R8A or E9P, previously reported to abolish BoxB binding in biochemical assays (31, 34), did not significantly affect EV titers. The λN domain is arginine-rich, and mutation of any one of its five arginine residues disrupts BoxB binding (34). NMR analyses of the λN:BoxB complex confirmed that R6, R7, R8, R10, and R11 directly contact BoxB (32). A construct carrying alanine substitutions at all five sites (R6/7/8/10/11A) yielded EVs at 22,000 DU/mL, a significant reduction relative to the positive control but still higher than the non-VSV-G control (Fig. 2F). These results demonstrate that the λN domain within EVPC-5 is essential for efficient Cre mRNA packaging. The residual EV production likely reflects passive mRNA loading. Compared with the no-GZR condition (SI Appendix, Fig. S1D), where neither λN nor EABR is fused to the chimeric protein and titers reached only at 1,800 DU/mL, the ΔλN variant retains the C-terminal EABR domain that promotes EV biogenesis. Thus, although both conditions lack λN and load mRNA passively, the ΔλN construct releases more EVs. As observed for the PH domain, extensive mutations are required to abolish λN function, likely because I3-01-mediated oligomerization increases the avidity of λN variants for their RNA-binding partner.

#### Deletion of the BoxB element

Deletion of the BoxB element in the 3′ UTR of the Cre mRNA further supports the hypothesis of multivalent interactions between BoxB-containing RNA and λN. The BoxB element forms a 19-nucleotide hairpin with a 5-nucleotide loop (GAAGA or GAAAA) and a 7–base-pair stem that stabilizes the loop in the conformation required for high-affinity λN binding. The λN peptide binds tightly to the wild-type BoxB hairpin but only weakly to RNAs carrying mutations in the loop or stem (31). NMR analyses of the λN:BoxB complex further revealed that the GNRA fold (N = A/C/T/G; R = A/G), where G is the first and A the fifth nucleotide in the BoxB loop, is essential for λN recognition (32). However, deletion of the BoxB element in EVPC-5 reduced EV titers by 3.5-fold, which was not statistically significant relative to the positive control (*p* = 0.12) (Fig. 2A). This remaining packaging likely reflects the increased avidity of λN conferred by I3-01-mediated oligomerization, allowing it to bind BoxB-like motifs within the Cre mRNA even with low intrinsic affinity, thereby facilitating mRNA loading in the absence of canonical BoxB hairpins.

#### Deletion or loss-of-function variants of the EABR domain

Deletion of the EABR domain reduced EV titers 10-fold but remained significantly higher than the non-VSV-G control (Fig. 2A). The EABR domain of CEP55 mediates interactions with both ALIX and the ESCRT-I subunit TSG101. Critical residues have been identified: single-point mutations W184A, Y187A, or R191A abolish detectable binding to ALIX and likely also to TSG101, while E192A substantially reduces affinity for ALIX (20). A construct carrying W184A/Y187A double mutations produced EVs at a 3-fold lower titer than EVPC-5, which was not statistically significant relative to the positive control (*p* = 0.10). Addition of R191A and E192A mutations to generate a W184A/Y187A/R191A/E192A quadruple mutant reduced EV titers by 6-fold, a statistically significant decrease, though still higher than the non-VSV-G control. To further evaluate ESCRT involvement in EV production, a dominant-negative VPS4A mutant (E228Q) was coexpressed with EVPC-5 and VSV-G (11, 16, 52). This combination reduced EV titers by 100-fold relative to the positive control but remained significantly higher than the non-VSV-G group. Because VPS4 is a central ATPase driving ESCRT-dependent membrane remodeling (53, 54), these results indicate that EV production by EVPC-5 requires the ESCRT pathway and that other regions beyond EABR can recruit ESCRT machinery to support EV release. An ESCRT-independent mechanism also contributes to EV production. Moreover, Positioning the EABR domain at the C terminus of I3-01 may enhance the avidity of EABR for ALIX and TSG101, necessitating more extensive mutations to fully disrupt ESCRT recruitment in this context (Fig. 2G).

Overall, these results show that only deletion or loss-of-function mutations in the membrane-targeting PLCδ1-PH domain reduced EV titers to levels not significantly different from the non-VSV-G control. Mutations in other components either partially reduced EV titers (HCV NS3/4A protease, λN, and the EABR domain) or had no significant effect (I3-01 nanocage and BoxB).

### Maximizing EV mRNA delivery

The small size of λN allowed multiple tandem copies to be included using a 16–amino-acid GS linker (SGGGG)_3_S to connect them, with each copy adding only 111 nucleotides to the ∼5-kb Cre mRNA and 37 amino acids to the ∼700-aa chimeric protein. Constructs containing 1, 3, 5, 7, or 9 tandem λN copies (EVPC-5 to EVPC-9; Fig. 3A) produced EVs at titers that varied nonlinearly with λN valency. The 5-copy construct (EVPC-7) achieved the highest yield, a 12-fold increase compared with the 1-copy construct (EVPC-5). Increasing the λN copy number to three (EVPC-6) enhanced EV production sixfold, whereas seven copies (EVPC-8) yielded a fivefold increase. However, further increasing λN repeats to nine (EVPC-9) reduced EV production to a level comparable to the 1-copy construct (Fig. 3B). Given that λN contains five positively charged arginine residues, the reduction in EV production at higher λN copy numbers may reflect competition between λN binding to Cre-containing mRNA and electrostatic interactions with negatively charged cellular membranes (55), which would be amplified by multivalent λN display on the I3-01 scaffold.

**Fig. 3.**
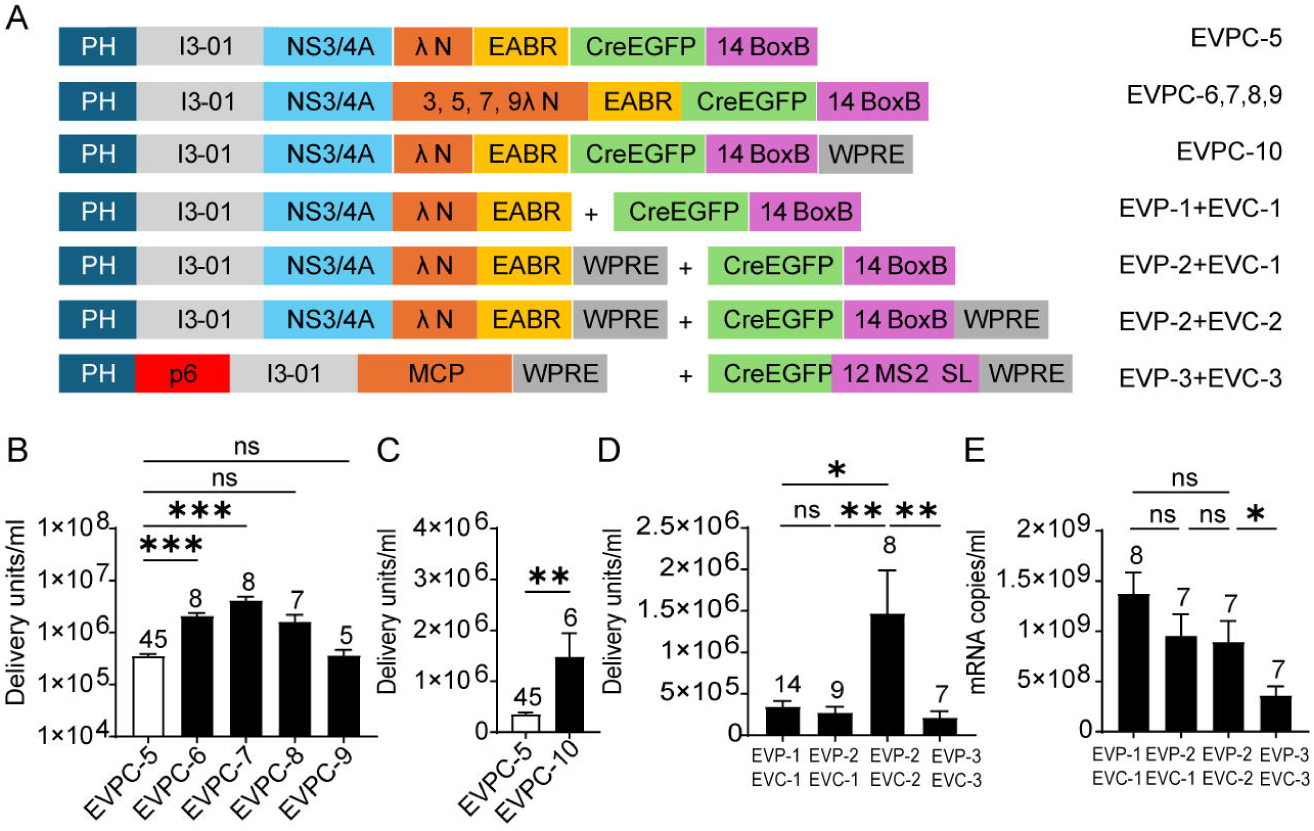
Optimization of λN copy number and WPRE placement in the EVPC 3′-UTR. (A) Design schematic. (B) EV functional titers for EVPC variants encoding 1, 3, 5, 7, or 9 copies of λN. *p*-values were analyzed using the Kruskal–Wallis test followed by Dunn’s multiple comparisons test vs. EVPC-5 control. (C) EV functional titers for EVPC-10 with WPRE in the 3′ UTR. *p*-value was calculated using Mann–Whitney test. (D) Comparison of EV functional titers for various EVP/EVC constructs. *p*-values across the first three groups were calculated using the Kruskal–Wallis test followed by Dunn’s multiple comparisons test. The comparison between the third and fourth groups was evaluated using Mann–Whitney test. (E) mRNA export control. RT-qPCR quantification of Cre mRNA in producer-cell supernatants for the four groups in (D). *p*-values across the first three groups were calculated using Tukey’s multiple comparisons test following ordinary one-way ANOVA. The comparison between the third and fourth groups was evaluated using an unpaired t-test.

Addition of the woodchuck hepatitis virus posttranscriptional regulatory element (WPRE) to the 3′ UTR downstream of the BoxB motif (EVPC-10) significantly increased EV titers compared with the construct lacking WPRE (EVPC-5; Fig. 3C). To investigate whether WPRE enhances EV titers by increasing nuclear export of Cre-containing mRNA—thereby facilitating greater loading into EVs in producer cells—or by enhancing mRNA stability and promoting higher levels of Cre protein translation in recipient cells (56–58), the EVPC-5 plasmid was divided into separate EV-packaging (EVP) and EV-cargo (EVC) vectors (Fig. 3A). Cotransfection of these separated constructs with VSV-G produced EVs at titers comparable to the single-vector control. Addition of WPRE to the packaging vector (EVP-2) had no effect, whereas its inclusion in the cargo vector (EVC-2) significantly increased EV titers relative to the no WPRE control (Fig. 3D). RT-qPCR analysis of producer-cell supernatants revealed no significant differences in exported Cre mRNA levels across groups (Fig. 3E), suggesting that the WPRE primarily enhances post-entry mRNA stability or translation in recipient cells rather than EV loading in producer cells. For comparison with the EPN24-MCP RNA delivery system described by Horns et al., we generated EVP-3 and EVC-3 plasmids. EVP-3 encodes EPN24-MCP using the same vector backbone as EVP-2, and EVC-3 is analogous to EVC-2 but encodes CreEGFP mRNA containing 12 MS2 stem-loop repeats, matching the copy number used previously (11). Coexpression of the MCP construct (EVP-3) and the MS2 stem loop construct (EVC-3) with the VSV-G plasmid yielded EVs with titers approximately sevenfold lower than those from the λN (EVP-2) + BoxB construct (EVC-2) combination, and mRNA export levels were about 2.5-fold lower; both reductions were statistically significant (Fig. 3D and E).

### Multiple Copies of λN Enable BoxB-Independent RNA Loading and Functional Delivery

Based on the λN copy number titration and the WPRE results, we generated EVP-4, containing five λN copies (Fig. 4A). Although the WPRE in the packaging vector was dispensable, it was retained to avoid introducing additional variables in subsequent comparisons. Coexpression of the packaging plasmid EVP-4, the cargo plasmid EVC-2 containing 14 BoxB copies, and VSV-G produced EVs with a sixfold higher titer compared to EVP-2, which contains a single λN domain. Deletion of the BoxB element in the cargo RNA (EVC-4) produced EVs at titers not significantly different from the BoxB-containing control (EVC-2). RT-qPCR analysis revealed no change in exported mRNA levels. Thus, in EVP-4, which contains five λN repeats, BoxB sequences were dispensable for efficient mRNA packaging and functional EV production (Fig. 4 B–D).

**Fig. 4.**
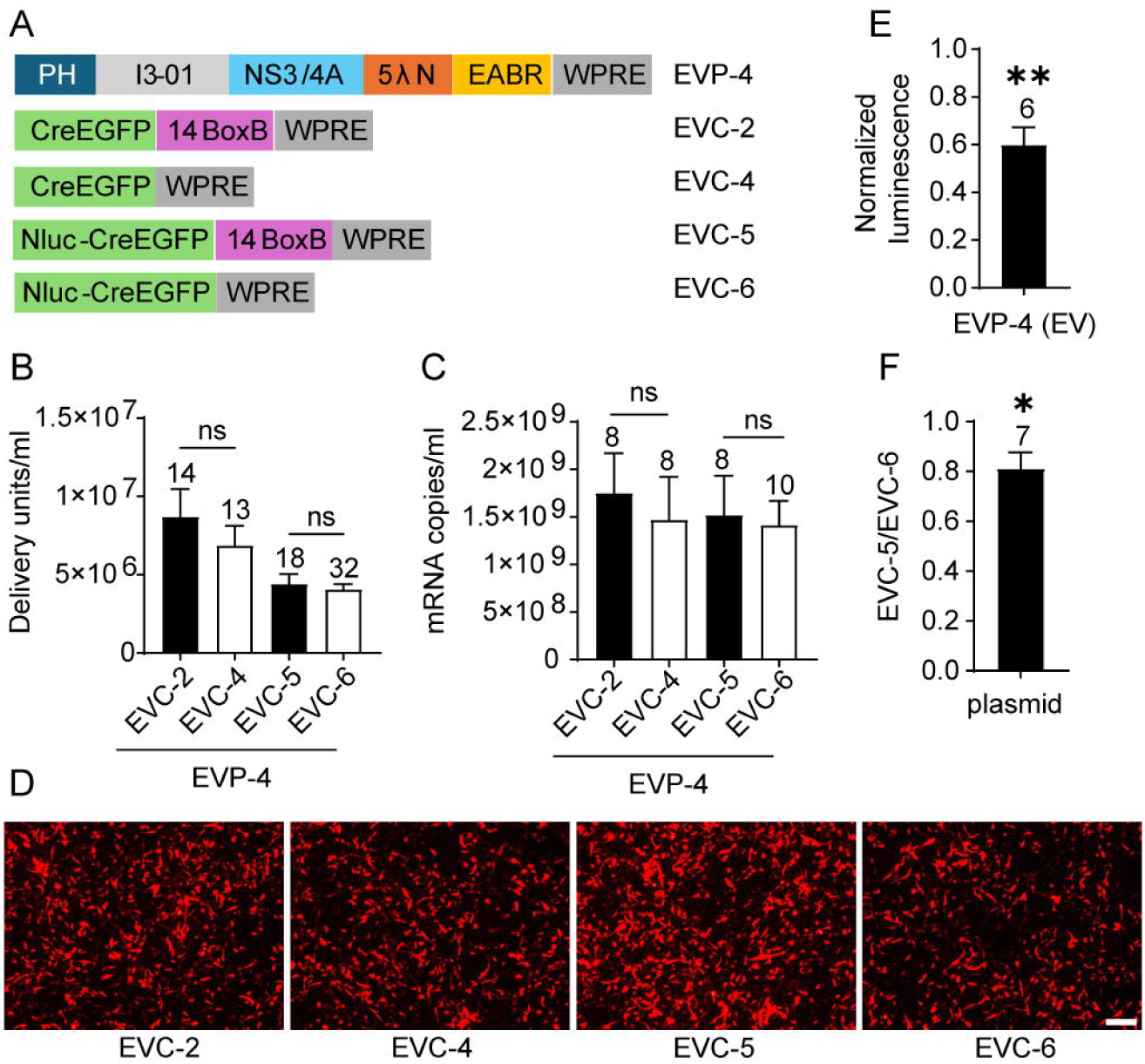
Test of BoxB requirement in EVP-4–mediated RNA export and EV functional titers. (A) Design schematic. (B) In combination with EVP-4, EV functional titers of EVCs with or without canonical BoxB elements. EVC-2 vs EVC-4, unpaired t-test; EVC-5 vs EVC-6, Mann–Whitney test. (C) mRNA levels in supernatants from the four groups shown in (B). EVC-2 vs EVC-4, unpaired t-test; EVC-5 vs EVC-6, unpaired t-test. (D) Representative fluorescence images of DsRed⁺ BHK-21 recipient cells from the first dilution well of the EV titration assay for the four groups in (B). Scale bar, 100 μm. (E) In 293T recipient cells exposed to EVs, NanoLuc activity for EVC-5 normalized to EVC-6. The *p*-value was calculated using a one-sample t-test with a theoretical mean of 1. (F) Direct-transfection control. EVC-5 and EVC-6 plasmids were transfected into 293T cells, i.e. not packaged in EVs. The EVC-5/EVC-6 NanoLuc ratio (fold change) was computed within batches and compared with 1 by a one-sample t-test.

To assess whether BoxB influences translation of delivered mRNA, NanoLuc-based cargo plasmids were generated: EVC-5 (14× BoxB) and EVC-6 (no BoxB). When coexpressed with the packaging plasmid, EVP-4 (5 λN copies), and VSV-G, both produced comparable EV titers and similar levels of mRNA export (Fig. 4 B–D). These results reinforce the conclusion that BoxB is not needed on the cargo RNA when a packaging construct with multiple λN copies is used. EVC-6 alone led to the export of 3.1 × 10□ mRNA copies/mL (n = 4), a ∼450-fold decrease compared to the group that included the packaging plasmid EVP-4 and VSV-G. Because VSV-G expression did not affect mRNA export (SI Appendix, Fig. S1C), this result indicates that the packaging construct EVP-4 accounts for the bulk of measurable mRNA export. To assess delivery and translation of cargo mRNA, NanoLuc luciferase assays were performed. NanoLuc activity in the EVC-5 (14× BoxB) group was normalized to that of EVC-6 group (no BoxB). Despite similar levels of mRNA export, BoxB-containing EVs (EVC-5) produced only 60% of the luminescence of BoxB-deleted EVs (EVC-6), a slight but significant reduction (Fig. 4E). Because RNA hairpins in the 3’UTR can influence mRNA translation and stability (59), we compared expression from plasmid transfection of EVC-5 and EVC-6 in 293T cells in the absence of the packaging plasmid. EVC-5 resulted in 80% of the luminescence of EVC-6 (Fig. 4F), indicating that multiple BoxB repeats suppress mRNA translation independently of the packaging chimera. Thus, removal of BoxB sequences enhances protein translation without affecting RNA export.

### BoxB-Independent mRNA Packaging and Delivery Achieved Through Multivalent λN Avidity

Because exported mRNA levels were similar for BoxB-containing and BoxB-deleted cargos, we hypothesized that the I3-01 nanocage in EVP-4 provides a high effective λN valency per vesicle. This increased avidity would allow weak BoxB-like motifs to mediate RNA loading even in the absence of canonical BoxB sequences. To test this, we generated packaging constructs with reduced λN valency. Constructs containing trimeric KDPG aldolases from *T. maritima* (EVP-5; PDB 1WA3) and *E. coli* (EVP-6; PDB 2C0A) were compared with EVP-4. These trimeric packaging constructs lack the I3-01 nanocage assembly of EVP-4 and therefore reduce λN valency by 20-fold. EVP-2, which contains only a single λN peptide on the I3-01 scaffold, results in a 5-fold reduction in valency, while EVP-7, which combines the trimeric *T. maritima* KDPG aldolase with a single λN peptide, leads to a 100-fold reduction in λN valency compared with EVP-4 (Fig. 5A). When paired with BoxB-containing cargo (EVC-5), the trimeric constructs EVP-5 and EVP-6 produced EV titers and mRNA export levels comparable to EVP-4, whereas the I3-1 nanocage based single λN containing EVP-2 yielded significantly reduced titers but retained comparable mRNA export. In contrast, when BoxB was removed from the EVC5 cargo (forming EVC-6), both EVP-2 and EVP-5 showed significant reductions in EV production and mRNA export relative to EVP-4, while trimeric E. coli KDPG aldolase containing EVP-6 exhibited only a minor, non-significant decrease (Fig. 5B and C). EVP-7, which has the lowest λN valency among all constructs, showed the most pronounced reduction in both titers and RNA export compared to the EVP-4 control (Fig. 5D and E). Compared with EVP-4, which has the highest λN valency and maintained similar EV titers and mRNA export with or without BoxB (EVC-5 vs. EVC-6), constructs with reduced λN valency (EVP-2, EVP-5, and EVP-6) displayed only modest BoxB dependence, whereas EVP-7 showed the strongest dependence. Together, these results demonstrate that λN valency is a key determinant of BoxB-independent RNA packaging, with EVP-4 uniquely maintaining high packaging efficiency even in the absence of canonical BoxB motifs.

**Fig. 5.**
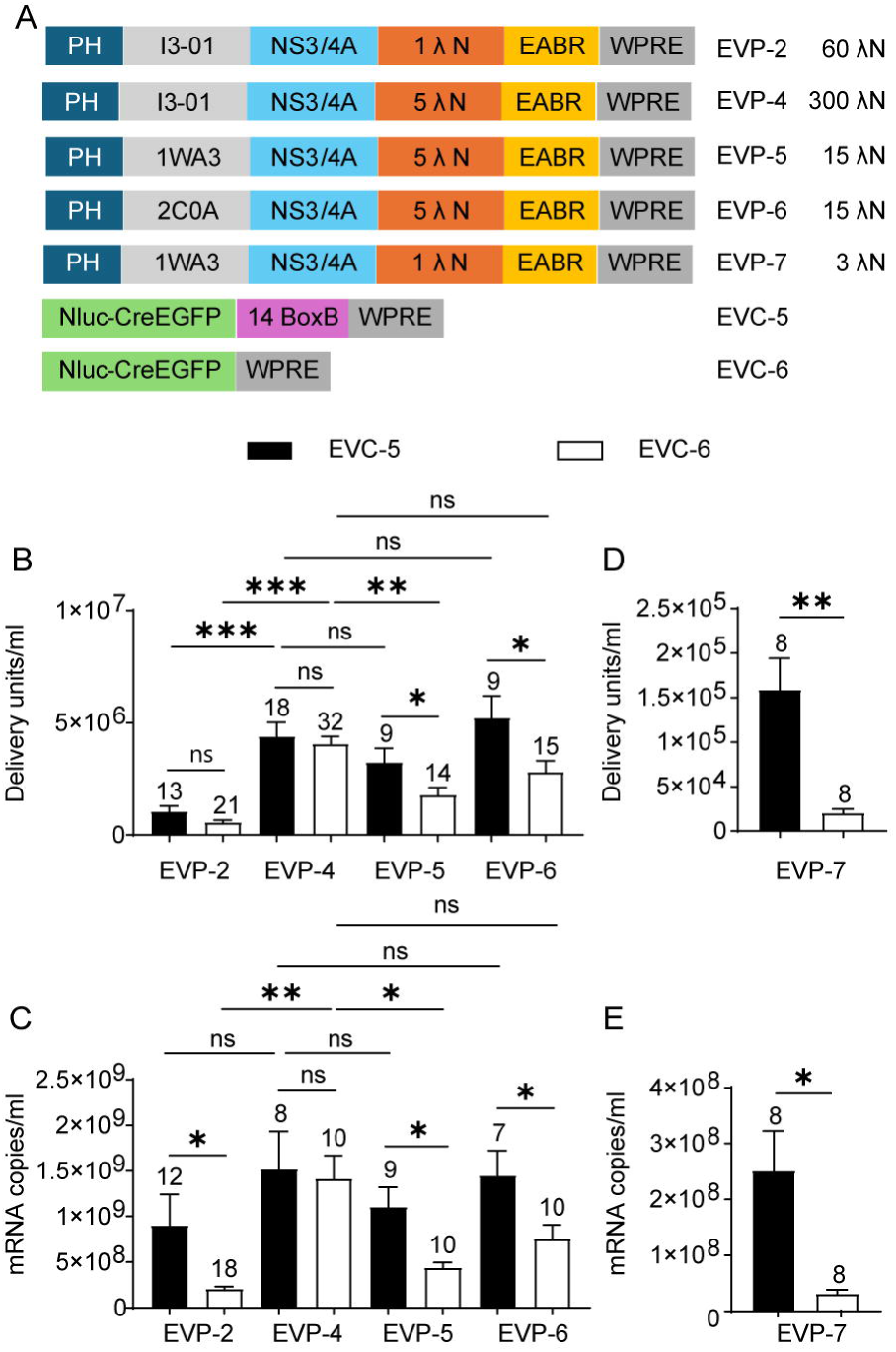
Effect of λN valency on BoxB-dependent mRNA loading and functional delivery. (A) Design schematic, with the number of λN copies per EV indicated on the right. (B) EV functional titers of EVP variants in combination with EVC-5 or EVC-6. *p* values across the four EVP groups paired with EVC-5 or EVC-6 were calculated using Kruskal–Wallis test followed by Dunn’s multiple-comparison test versus the EVP-4 group. Comparisons between EVC-5 and EVC-6 paired with the same EVP were performed using an unpaired t test (EVP-5) or the Mann–Whitney test (EVP-2, EVP-4, and EVP-6). (C) mRNA export levels for the eight groups shown in (B). *p*-values across the four EVP groups paired with EVC-5 were calculated using the Kruskal–Wallis test followed by Dunn’s multiple-comparison test versus the EVP-4 group. *p*-values across the four EVP groups paired with EVC-6 were calculated using Welch’s ANOVA followed by Dunnett’s T3 multiple-comparison test versus the EVP-4 group. Comparisons between EVC-5 and EVC-6 paired with the same EVP were performed using an unpaired t test (EVP-4 and EVP-5), Welch’s t test (EVP-6), or the Mann–Whitney test (EVP-2). (D) EV functional titers of EVP-7 in combination with EVC-5 or EVC-6. The *p*-value was calculated using a paired t test. (E) mRNA export levels for the two groups shown in (D). The *p*-value was calculated using a paired t test.

To examine whether putative BoxB-like motifs contribute to RNA loading, we first analyzed the EVC-4 cargo, which encodes CreEGFP-WPRE and lacks the 14 canonical BoxB repeats present in EVC-2. A search for the consensus λN-binding motif (UGNANAA, N = A/C/T/G) (31, 32) within EVC-4 identified five candidate sites—one each in EGFP and WPRE and three within the Cre coding region. To dissect their contributions, we generated sequential derivatives of EVC-4. In EVC-7, the EGFP sequence was deleted. In EVC-8, the WPRE was removed, and point mutations were introduced at the G8 position of two of the three BoxB-like motifs within Cre, while the third motif was left unmodified to preserve the encoded amino acids (SI Appendix, Fig. S2 A). These modifications effectively depleted the endogenous BoxB-like motifs present in EVC-4. When co-expressed with the nanocage-containing packaging construct EVP-4, EVC-8 exhibited a statistically significant threefold reduction in exported mRNA relative to the BoxB-containing construct EVC-2. Restoring 14 canonical BoxB repeats to the 3’ UTR of EVC-8 generated EVC-9, which recovered mRNA export to levels comparable to EVC-2 (SI Appendix, Fig. S2C). Because WPRE influences EV titers, we compared only EVC-8 and EVC-9, both lacking WPRE, for functional EV production. Incorporation of 14× BoxB in EVC-9 resulted in a 2.2-fold increase in EV titers relative to EVC-8 (SI Appendix, Fig. S2D). These findings suggest that one or more BoxB-like motifs in EVC-4 facilitate RNA packaging, likely through the formation of **G**N**RA** folds by the λN-binding consensus sequence (U**G**N**A**N**A**A), which are stabilized by flanking nucleotides (32). Overall, tandem λN repeats and the self-assembling nanocage enhance λN valency and RNA avidity, rendering canonical BoxB sequences dispensable for mRNA packaging and functional EV production.

### A Human Trimeric Scaffold (CLYBL) Can Substitute for the Bacterial Nanocage and Removal of HCV Protease Preserves Functional EV Production

Based on the finding that the I3-01 nanocage could be replaced by trimeric bacterial KDPG aldolases without causing a substantial loss in titer, we next examined whether a human structural homolog could substitute for these bacterial scaffolds. This may improve translational potential by reducing immunogenicity. A structural similarity search on the NCBI VAST+ website using the *T. maritima* KDPG aldolase (PDB 1WA3) as the input identified human CLYBL (PDB 5VXO) as the only eukaryotic protein with a complete structural match (Fig. 6A). CLYBL encodes a 340–amino-acid mitochondrial enzyme expressed in most tissues (60). To localize it to the cytoplasm and prevent metabolic interference, we deleted the N-terminal mitochondrial targeting sequence (residues 1–22) and introduced the catalytically inactive D320A mutation (61). Replacing the I3-01 nanocage in the packaging plasmid EVP-4 (five λN repeats, I3-01 nanocage, NS3/4A wild-type protease) with this human variant generated EVP-8 (five λN repeats, human CLYBL scaffold, NS3/4A wild-type protease) (Fig. 6B).

**Fig. 6.**
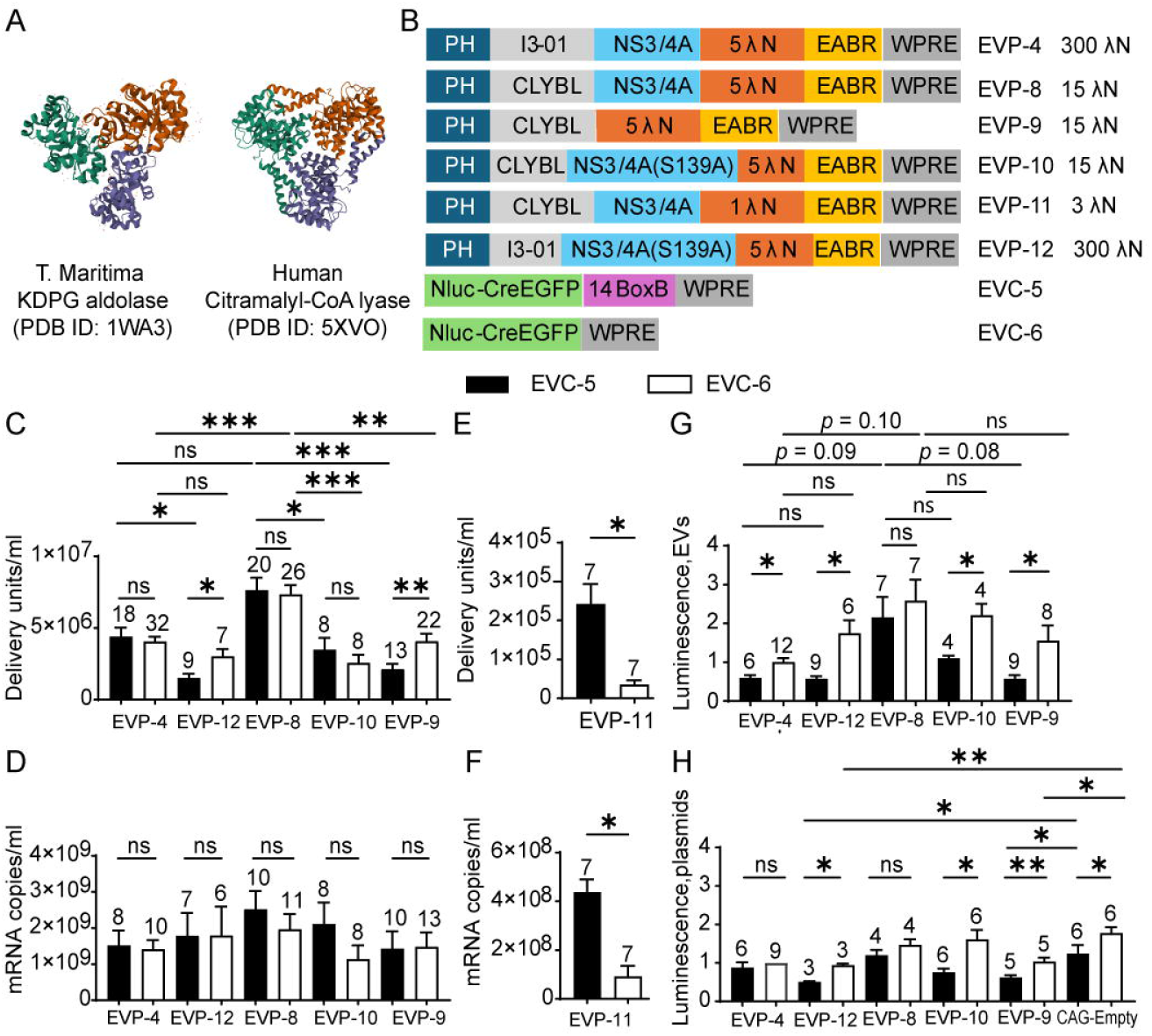
Test of a human trimeric protein (CLYBL) as a substitute for the bacterial nanocage and the effect of removal of HCV NS3/4A. (A) VAST+ search using *T. maritima* KDPG aldolase (PDB 1WA3) identified human citramalyl-CoA lyase (PDB 5VXO) as the only complete structural match among Eukaryota. (B) Design schematics. (C) EV functional titers for EVP variants. Each was tested with EVC-5 and EVC-6. *p*-values across the five EVP groups paired with EVC-5 or EVC-6 were calculated using Kruskal–Wallis test followed by Dunn’s multiple comparisons test (for EVC-5) or Welch’s ANOVA followed by Dunnett’s T3 multiple comparisons test (for EVC-6). Comparisons between EVC-5 and EVC-6 paired with the same EVP were performed using an unpaired t-test (EVP-8 and EVP-12), Welch’s t test (EVP-9), or Mann–Whitney test (EVP-4 and EVP-10). (D) mRNA export levels of the ten groups shown in (C). *p*-values across the five EVP groups paired with EVC-5 or EVC-6 were calculated using Kruskal–Wallis test. As the overall test did not reveal statistically significant differences, no post-hoc multiple comparisons were conducted. Comparisons between EVC-5 and EVC-6 paired with the same EVP were performed using an unpaired t-test (EVP4, EVP-8, and EVP-12) or Mann–Whitney test (EVP-9 and EVP-10). (E) EV functional titers of EVP-11 in combination with EVC-5 or EVC-6. *p*-values were calculated using the Wilcoxon test. (F) mRNA export levels of the two groups shown in (E). *p*-value was calculated using the Wilcoxon test. (G) EV post-delivery translation activity. NanoLuc activities were normalized to EVP-4 + EVC-6. *p*-values across the five EVP groups paired with EVC-5 or EVC-6 were calculated using Welch ANOVA followed by Dunnett’s T3 multiple comparisons test. Comparisons between EVC-5 and EVC-6 paired with the same EVP were performed using an unpaired t-test (EVP-4 and EVP-8) or Welch’s t test (EVP-9, EVP-10, and EVP-12). (H) Direct-transfection control. In the absence of GZR, 293T cells were directly transfected with EVPs bearing active protease (EVP-4, EVP-8) or no/inactive protease (EVP-9, EVP-10 [S139A], EVP-12 [S139A]), together with EVC-5 or EVC-6. A CAG-empty control was included to match total plasmid mass. NanoLuc activity was measured and normalized to the EVP-4 + EVC-6 condition. With the exception of the EVP-4 + EVC-6 group, comparisons between the CAG-empty control and EVP groups paired with either EVC-5 or EVC-6 were evaluated using ordinary one-way ANOVA followed by Dunnett’s multiple comparisons test (for EVC-5) or the Kruskal–Wallis test followed by Dunn’s multiple comparisons test (for EVC-6), relative to the CAG-empty control. Comparison between EVC-5 and EVC-6 paired with the same EVP were performed using a paired t-test (EVP8, EVP-9, EVP-12, and CAG-empty control) or Wilcoxon test (EVP-10). Comparisons of EVP-4 paired with EVC-5 or EVC-6 were performed using a one-sample t-test against a theoretical mean of 1.

When paired with either BoxB-containing (EVC-5) or BoxB-deleted (EVC-6) cargo, the CLYBL construct (EVP-8) produced modestly higher EV titers (∼2-fold) than the I3-01 nanocage construct (EVP-4), although this increase reached statistical significance only with BoxB-deleted cargo (EVC-6). mRNA exports from the CLYBL-producer cells (EVP-8) showed a nonsignificant upward trend relative to the I3-01 nanocage producer cells (EVP-4; Fig. 6C and D). Like the I3-01 nanocage, the CLYBL scaffold showed no significant difference in EV titer and mRNA export between BoxB-containing and BoxB-deleted cargos, indicating that CLYBL can also support BoxB-independent RNA packaging and functional EV production.

To assess whether λN valency remains critical within the CLYBL scaffold, we reduced the number of λN repeats from five in EVP-8 to one in EVP-11. This single-λN construct exhibited markedly reduced EV titers and mRNA export relative to the 5-λN CLYBL construct, for both BoxB-containing (EVC-5) and BoxB-deleted (EVC-6) cargos. EVP-11 also showed a pronounced dependence on BoxB, with significantly lower output for the BoxB-deleted cargo, mirroring the behavior of EVP-7 (single λN on the trimeric KDPG aldolase scaffold). These results indicate that λN valency, rather than scaffold identity, is the primary determinant of RNA packaging efficiency (Fig. 6E and F).

To further humanize the system, we deleted the HCV NS3/4A protease from the CLYBL construct, to create EVP-9. To test the effect of protease removal vs. lack of protease enzymatic activity, we also created EVP-10, which has the protease inactive allele, S139A. These two constructs were compared to a CLYBL construct with the wild type protease (EVP-8) and with nanocage-based controls encoding the I3-01 scaffold and active protease (EVP-4), or the I3-01 scaffold and the protease inactive mutant (EVP-12) (Fig. 6B). When paired with either BoxB-containing (EVC-5) or BoxB-deleted (EVC-6) cargo, removal or inactivation of the NS3/4A protease in the CLYBL scaffold caused significant reductions in EV titers relative to the wild-type protease construct (EVP-8). Similar trends were observed for the nanocage scaffold, where EVP-12 (S139A) produced a significant decrease with BoxB-containing cargo (EVC-5) but not with BoxB-deleted cargo (EVC-6). Although differences in functional EV titers were observed (Fig. 6C), RT-qPCR analysis revealed no significant changes in exported mRNA levels between BoxB-containing (EVC-5) and BoxB-deleted (EVC-6) cargos packaged by these constructs (Fig. 6D). Moreover, neither protease removal nor inactivation significantly affected total RNA export in either the CLYBL or I3-01 scaffolds, indicating that protease activity modulates functional delivery rather than RNA packaging per se.

To assess RNA translation after delivery, NanoLuc luciferase assays were performed following delivery of EVs into recipient cells. The CYBYL construct with active protease (EVP-8) produced the highest luminescence in average among all constructs, regardless of whether the RNA cargo had BoxB. However, these increases did not reach statistical significance when compared to other constructs. This result indicates that CLYBL with an active protease supports efficient functional RNA delivery and translation. Within each EVP group, BoxB-containing cargo (EVC-5) consistently yielded lower translation activity than BoxB-deleted cargo (EVC-6), and this difference was statistically significant for all constructs except EVP-8. These findings identify the CLYBL construct with active protease (EVP-8) as the most efficient construct, yielding the highest functional titers and translational activity so far (Fig. 6G).

To determine whether protease-mediated cleavage influences mRNA translation, 293T cells were co-transfected with EVPs and either BoxB-containing (EVC-5) or BoxB-deleted (EVC-6) cargo RNAs in the absence of protease inhibitor. NanoLuc assays showed that constructs with functional protease (EVP-4 and EVP-8), which are expected to cleave the λN–EABR fragment from the packaging protein, exhibited no significant difference in translation between BoxB-containing (EVC-5) and BoxB-deleted (EVC-6) cargo. By contrast, constructs lacking protease activity (EVP-9, EVP-10, and EVP-12), which are expected to retain the full-length chimeric packaging protein, showed significantly lower translation when paired with the BoxB-containing cargo (EVC-5). Furthermore, in the absence of protease activity, the protease negative constructs (EVP-9 and EVP-12) suppressed translation significantly below empty-vector levels, suggesting that accumulation of the uncleaved chimeric protein inhibits mRNA translation (Fig. 6H).

Overall, replacing the bacterial nanocage with the human CLYBL scaffold maintained (EVP-9) or enhanced (EVP-8) EV production, and removal of the viral protease further humanized the design without compromising function. The most humanized EVP-9 construct—retaining only the bacteriophage λN peptide while lacking both bacterial and HCV-derived structural components—efficiently packaged and delivered BoxB-deleted EVC-6 mRNA, achieving high functional titers and translational output.

### ENTH-based Membrane Targeting and λN Valency Optimization enhance EV Titers, mRNA Export, and Translation

Having established the construct with CLYBL without protease as a strong candidate for EV-based RNA delivery with improved translational activity, we next optimized its components and evaluated performance across three key parameters: EV titers, mRNA export, and translational activity of EV-delivered mRNA. This optimization was also intended to clarify mechanisms underlying efficient EV production.

Phosphatidylinositol-4,5-bisphosphate [PI(4,5)P₂] is enriched at the plasma membrane (62). To use this localization as a means to improve membrane targeting of the chimera and perhaps EV production, we tested whether a membrane targeting domain that showed higher specificity for PI(4,5)P₂ might lead to higher EV production. To this end, we replaced the rat PLCδ1 PH domain with the ENTH domain (residues 1–158) of human epsin-1. Epsin-1 is an endocytic adaptor protein whose ENTH domain exhibits high affinity for PI(4,5)P₂ and has been used as a biosensor for this lipid (63). To further enhance specificity, we introduced the S4W mutation into the ENTH domain, which increases PI(4,5)P₂ binding while reducing affinity for PI(3,4)P₂ (63), a lipid enriched in both plasma membrane and early endosomes (62). Because VSV-G is synthesized in the ER and trafficked via the Golgi to the plasma membrane, with only occasional localization to endosomal compartments (64–68), enhanced plasma-membrane targeting was expected to promote colocalization with VSV-G, the efficient fusogen. The packaging construct with ENTH S4W, CLYBL, and 5 λN (EVP-14) was generated by replacing the PH domain in the packaging construct EVP-9, which has CLYBL and the protease deletion. Two additional variants were created containing seven tandem λN repeats: EVP-13 (PH, CLYBL, 7 λN) and EVP-15 (ENTH S4W, CLYBL, 7 λN). These constructs were tested for whether the λN copy number that we previously optimized for the I3-01 nanocage scaffold also was optimal for the human CLYBL scaffold (Fig. 7A).

**Fig. 7.**
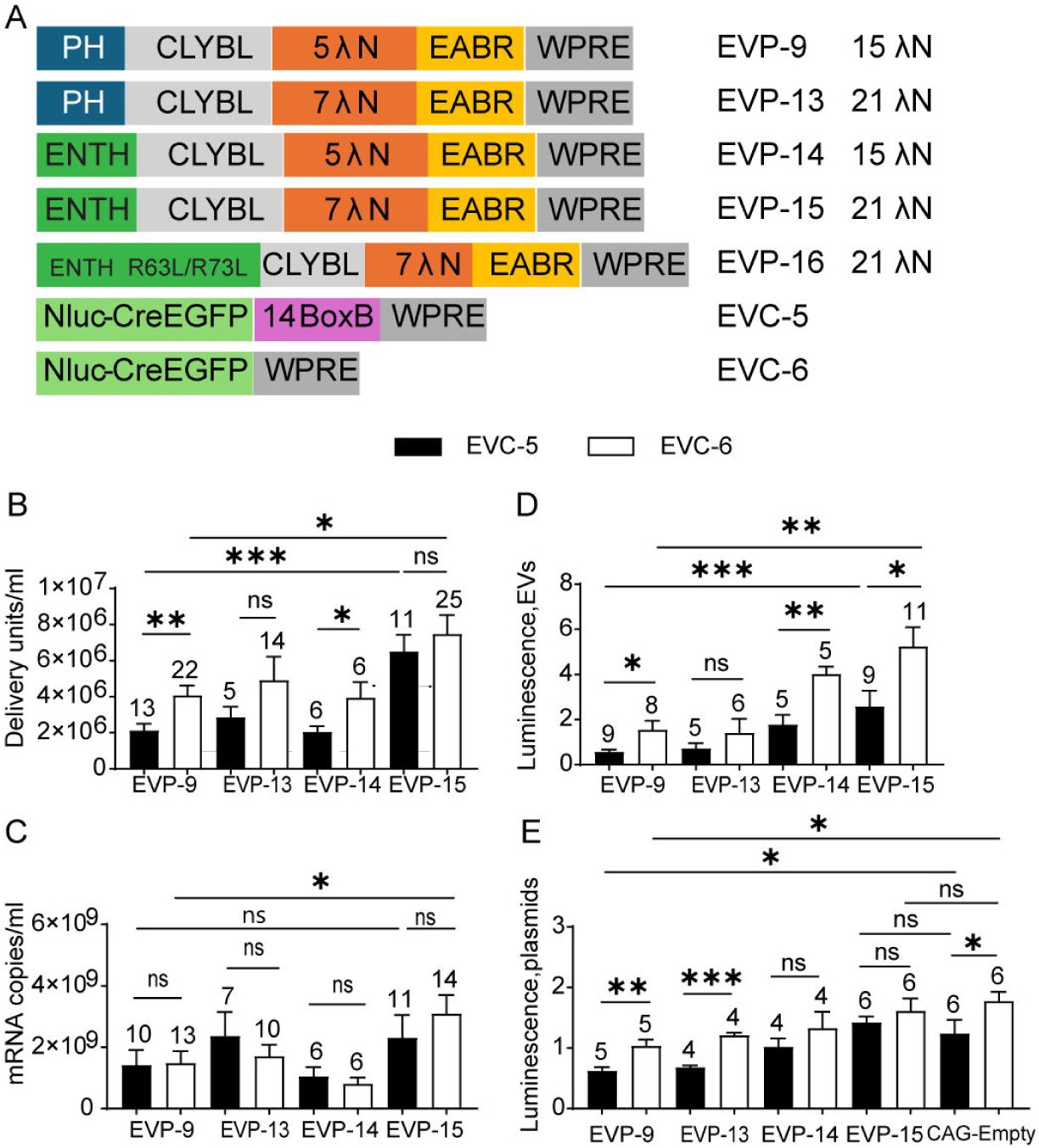
Test of human epsin1 ENTH as a substitute for rat PLCδ1-PH and effect of λN copy number. (A) Design schematics. (B) EV functional titers for EVP variants. Each was tested with EVC-5 and EVC-6. *p*-values across the first four EVP groups paired with EVC-5 or EVC-6 were calculated using Kruskal–Wallis test followed by Dunn’s multiple comparisons test. Comparisons between EVC-5 and EVC-6 paired with the same EVP were performed using a paired t-test (EVP-14), Welch’s t test (EVP-9), or Mann–Whitney test (EVP-13 and EVP-15). (C) mRNA export levels of the ten groups shown in (B). *p*-values across the first four groups paired with EVC-5 or EVC-6 were calculated using Kruskal–Wallis test followed by Dunn’s multiple comparisons test (applied only to the EVC-6 groups). Comparisons between EVC-5 and EVC-6 paired with the same EVP were performed using an unpaired t-test (EVP-13), paired t-test (EVP-14), or Mann–Whitney test (EVP-9 and EVP-15). (D) EV post-delivery translation activity. NanoLuc activities were normalized to EVP-4 + EVC-6. *p*-values across the first four EVP groups paired with EVC-5 or EVC-6 were calculated using Kruskal–Wallis test followed by Dunn’s multiple comparisons test. Comparisons between EVC-5 and EVC-6 paired with the same EVP were performed using the Mann–Whitney test (EVP-13 and EVP-15) or Welch’s t test (EVP-9 and EVP-14). (E) Direct-expression control (no GZR). Cotransfection of EVPs with EVC-5 or EVC-6. A CAG-empty control was included to match total plasmid mass. NanoLuc activity was measured and normalized to the EVP-4 + EVC-6 condition. *p*-values for comparisons between the CAG-empty control and EVP groups paired with EVC-5 or EVC-6 were calculated using ordinary one-way ANOVA followed by Dunnett’s multiple comparisons test, relative to the CAG-empty control. Comparisons between EVC-5 and EVC-6 paired with the same EVP were calculated using a paired t-test.

When paired with either BoxB-deleted (EVC-6) or BoxB-containing cargo (EVC-5), the ENTH construct (EVP-15) produced significantly higher EV titers than the construct with the PH domain (EVP-9). Within each EVP group, constructs containing five λN repeats (EVP-9 and EVP-14), but not those with seven λN repeats (EVP-13 and EVP-15), showed significant differences between the BoxB-deleted (EVC-6) and BoxB-containing cargo (EVC-5) (Fig. 7B). To determine whether PI(4,5)P₂ binding is required, we generated EVP-16 (ENTH S4W R63L/H73L, CLYBL, 7 λN), in which the double mutation disrupts the PI(4,5)P₂-binding site of ENTH domain (69, 70). When paired with BoxB-deleted EVC-6, this PI(4,5)P₂-binding–defective construct produced only 4.5 × 10³ DU/mL EVs (n = 3), a 1,640-fold decrease compared with the ENTH S4W control construct (EVP-15), confirming that PI(4,5)P₂ binding is essential for EV production (SI Appendix, Fig. S3A). Exported mRNA levels measured by RT–qPCR was similar between the BoxB-deleted (EVC-6) and the BoxB-containing cargo (EVC-5) with each packaging construct, indicating that differences in functional titer were not due to gross changes in RNA release. However, with BoxB-deleted cargo (EVC-6), the construct with ENTH domain and seven λN repeats (EVP-15) exported significantly more mRNA than the construct with the PH domain and five λN repeats (EVP-9), consistent with improved mRNA packaging (Fig. 7C).

In NanoLuc luciferase assays following EV delivery, NanoLuc assays showed that the construct with the ENTH domain (EVP-15), combined with the BoxB-deleted cargo (EVC-6), produced the highest luminescence among CLYBL-based constructs. It exceeded both the PH domain CLYBL construct (EVP-9) and the nanocage control (EVP-4), indicating improved functional delivery and translation. Within the set of CLYBL constructs with the ENTH or the PH domain (EVP-9, EVP-14, and EVP-15), there were significant differences in the luminescence produced by the BoxB-deleted (EVC-6) and the BoxB-containing cargo (EVC-5) (Fig. 7D). To determine whether the EVP chimeric proteins directly influenced translation, NanoLuc assays were performed after cotransfection of EVPs with the EVC plasmids in the absence of protease inhibitor. With BoxB-deleted EVC-6, only the chimera with the PH domain (EVP-9) showed a significant reduction in NanoLuc activity relative to the CAG-empty control (Fig. 7E). In contrast, NanoLuc expression was restored to control levels across EVP-13–EVP-15, indicating that the translational inhibition caused by the EVP-9 chimeric protein was relieved. We hypothesize that reduced endosomal association of packaging protein chimeras facilitates rapid cytosolic release and alleviates translational drag (see Discussion; SI Appendix, Fig. S3C). Together with higher mRNA export, these results explain the maximal EV titers and enhanced translational output observed with the CLYBL construct with the ENTH domain (EVP-15).

Oligomers of >20 ENTH subunits can deform membranes and promote endocytosis, whereas smaller assemblies remain evenly distributed and do not induce membrane curvature (71). It was therefore of interest to examine the vesicle morphology of EVs derived from the ENTH domain construct (EVP-15). Negative-stain electron microscopy of EVs produced by EVP-15 + EVC-6 showed normal vesicle morphology with no evidence of membrane deformation, similar to EVs generated from PH domain–containing constructs (SI Appendix, Fig. S3B). These observations suggest that the ENTH domain in EVP-15-encoded chimeric protein likely assembles into oligomers smaller than 20-mers, which may be insufficient to drive the endocytic pathway. These smaller ENTH assemblies may favor membrane budding rather than endocytosis, supporting EV production.

Overall, replacing the membrane-targeting PH domain and increasing λN copy number in EVP-9 yielded EVP-15 (ENTH S4W, CLYBL, 7 λN), which achieved the highest EV titers, mRNA export, and translation efficiency in combination with BoxB-deleted cargo (EVC-6). Because EVP-15 encodes chimeric proteins composed almost entirely of human domains—epsin-1 ENTH, CLYBL, and CEP55 EABR—except for a 21-amino-acid λN peptide, they constitute a humanized EV-based mRNA delivery system projected to have reduced immunogenicity.

## Discussion

We present engineered EV constructs composed mainly of human-derived protein modules that achieve functional titers comparable to lentiviral vectors. By combining a biological titration assay to quantify functional titers with RT–qPCR of exported mRNA and post-delivery NanoLuc readouts, we demonstrate that optimization of membrane targeting, scaffold architecture, ESCRT recruitment, and λN copy number results in efficient mRNA delivery. In the titration assay, functional EV titers reflect the number of EVs that deliver sufficient Cre mRNA to produce detectable DsRed expression from multiple reporter plasmids. EVs that deliver Cre mRNA below this functional threshold are not counted in this assay, although their RNA content is still measured by RT-qPCR. EVs prepared without VSV-G serve as a negative control for the titration assay, accounting for plasmid or RNA carryover that is independent of fusogen-mediated delivery.

Our early constructs were based on EPN-24, which uses the I3-01 scaffold derived from the *T. maritima* KDPG aldolase and assembles into a 60-subunit nanocage (16, 44). Rather surprisingly, we found that nanocage formation is not required for functional EV production. Trimeric or even monomeric aldolase variants supported titers comparable to the designed 60-mer of I3-01, indicating that higher-order cage geometry is not the dominant driver of EV assembly or function. We speculate that this independence is in part due to the concentration of chimeric proteins within membrane microdomains, driven to these areas by membrane-binding domains in the chimera. Both the PLCδ1 PH domain and the epsin-1 ENTH domain bind PI(4,5)P₂ with high affinity, a lipid that forms microdomains on the inner leaflet of the plasma membrane (72). These membrane-binding domains may concentrate the chimeric proteins at the membrane, allowing EV assembly even in the absence of a nanocage. In addition, the human CEP55 EABR may contribute to assembly. It is a noncanonical coiled-coil domain that forms homodimers (20), and may promote further oligomerization of the chimeric proteins through higher-order coiled-coil interactions. Mutational analysis of the domains within the chimera further demonstrated that positioning these domains on the oligomeric scaffold increases the overall avidity of the PH, EABR, and λN domains, such that extensive mutations were required to disrupt function. By enhancing multivalent interactions, oligomerization can render weak-affinity interactions functionally comparable to strong-affinity binding partners, which explains why the EABR and p6 domains perform similarly in EV production despite their different intrinsic affinities for ESCRT components.

In contrast to the limited benefit of nanocage formation, λN valency emerged as the major determinant of functional EVs. Titration of λN copy number showed that multiple copies provide a significant increase in the functional titer, relative to only a single λN. Mechanistically, higher valency increases the probability that a single EV packages multiple RNA copies, shifting the per-EV mRNA distribution so that a larger fraction of vesicles exceeds the functional threshold for detectable recombination or translation in the titration assay. This reconciles the modest changes we observed in the total exported mRNA level with larger gains in functional titer. A striking result is that BoxB becomes dispensable with high λN valency: with five tandem λN copies arrayed on a multimeric scaffold, BoxB-deleted cargos are packaged and delivered at levels comparable to BoxB-containing controls. We hypothesize that multivalent λN–RNA contacts stabilize otherwise weak BoxB-like interactions within the cargo transcript. Consistent with this, reducing λN valency increases specificity for packaging and delivery of mRNAs bearing canonical BoxB hairpins. Conversely, tandem BoxB hairpins in the 3′ UTR modestly reduce translation, consistent with prior observations that structured 3′-UTR motifs can attenuate expression (59). Accordingly, the optimized mRNA-delivery construct uses EVC-6, which lacks tandem BoxB repeats in the 3′ UTR, for maximal expression efficiency.

Substituting the human trimeric CLYBL for the bacterial nanocage preserved or improved EV functional titers, mRNA export, and translation activity. Replacing the rat PLCδ1-PH with human epsin-1 ENTH (S4W), predicted to sharpen plasma-membrane specificity, also led to increased titers. Together with optimized λN valency, these changes restored or exceeded titers seen with nanocage- and HCV protease-containing designs. We speculate that after VSV-G–mediated fusion with the endosomal limiting membrane, ENTH-containing chimeras have little affinity for endosomal membranes, whereas PLCδ1-PH retains residual binding via PI(3,4)P₂, keeping a fraction of PH chimeras engaged with the endosome post-entry. In contrast, rapid cytosolic release of ENTH chimeras may favor dissociation of λN from weak BoxB-like sites and reduce translational drag. Consistent with this model, the humanized EVP-15, with ENTH (S4W) and without protease, exerts minimal inhibition on cargo translation, rendering protease cleavage dispensable, while maintaining high titers and translational output post-entry—simplifying manufacturing and lowering potential immunogenicity without compromising performance.

Compared with mRNA-based vaccines, which require only modest levels of antigen expression to elicit effective immune responses, mRNA therapeutics typically demand much higher and more sustained protein production to reach therapeutic levels (73). This modular EV system achieves efficient RNA delivery using VSV-G–mediated entry, a strategy shared with previous EV and VLP platforms, that improves endosomal escape (11–15). While VSV-G has been widely used to pseudotype lentiviral vectors owing to its broad tropism, particle stability, efficient cellular uptake, and endosomal escape, it is nevertheless susceptible to complement inactivation and can elicit adaptive immune responses that limit repeat dosing (74–76). Substituting VSV-G with other viral glycoproteins can improve serum resistance and yield serotypically distinct pseudotypes with potential for repeat in vivo dosing (77–80). For repeated administration, engineering humanized fusogens offers a rational route to mitigate immunogenicity. The biological titration assay described here provides a quantitative framework to evaluate such fusogen variants systematically.

In addition to VSV-G, the other nonhuman component in this modular EV system is the λN peptide derived from bacteriophage λ, which functions as an RNA-tethering motif. Unlike VSV-G, λN is a short (21-amino-acid) intra-vesicle peptide not exposed on the EV surface; therefore, the likelihood of triggering a humoral response after administration is reduced. Following cellular uptake, EV-associated λN peptides could, in theory, enter the MHC-I pathway through cross-presentation (81). However, because λN is a short peptide and is introduced via EV uptake rather than endogenous synthesis, only a small amount is likely to enter the cytosolic processing pathway, thereby limiting the diversity and abundance of MHC-I–restricted peptides. To assess the potential for cytotoxic T-cell activation, in silico immunogenicity analysis using NetMHCpan 4.2 across 12 HLA supertype representatives for 8–14-mer peptides predicted six weak binders and two stronger binders among 924 tested combinations (82, 83) (Dataset S1), indicating a very low likelihood of MHC class I–restricted T-cell activation. Although substitution of λN with several human RNA-binding domains was tested and yielded EVs at only ∼10□ DU/mL (data not shown), future iterations could adopt alternative human motifs if λN-specific immune responses are experimentally observed.

A complementary immunological question is whether humanized EV components might break self-tolerance. ENTH (from epsin-1), CLYBL, and EABR (from CEP55) are all widely expressed in different human tissues (60, 84). Following EV internalization and cross-presentation by recipient cells, if the abundance of these self-derived antigens is sufficient to engage self-reactive T cells that have escaped central tolerance, their recognition in the absence of inflammatory or co-stimulatory cues would be expected to reinforce peripheral tolerance rather than trigger immunity (85–89). However, VSV-G in viral particles engages the TLR4 signaling pathway in immune cells and induces proinflammatory cytokine secretion (90). In principle, such pro-inflammatory signals could facilitate cross-presentation of EV proteins on MHC-I and contribute to tolerance breakdown (87, 89, 91). Although the amount and surface density of VSV-G in EVs are likely lower than in authentic VSV particles and may not necessarily activate the TLR4 pathway, this possibility nonetheless warrants careful consideration for clinical translation. For single-dose therapies, non-humanized EVs may be sufficient to eliminate even theoretical risks of autoimmunity. For multi-dose regimens, replacing VSV-G with alternative viral glycoproteins that have not been reported to trigger TLR4 signaling (92, 93), or with human-derived fusogens, may reduce both anti-vector immunity and proinflammatory responses, thereby improving tolerability for multi-dose applications.

In summary, combined with a biological activity assay to quantify functional particles, this modular EV platform enables systematic optimization of membrane-targeting, scaffolding, ESCRT-recruiting, and RNA-tethering components to achieve efficient, human-compatible RNA delivery. Its flexible architecture supports rapid iteration and future engineering of EV fusogens and RNA-binding elements for enhanced efficacy and reduced immunogenicity.

## Abbreviations

CLYBL: citramalyl-CoA lyase beta-like protein
DU: delivery unit
EABR: ESCRT- and ALIX-binding region of CEP55
ENTH: epsin N-terminal homology
EPN: enveloped protein nanocage
ESCRT: endosomal sorting complex required for transport
EVC: EV cargo plasmid
EVP: EV packaging plasmid
EVPC: EV packaging-cargo plasmid
GZR: grazoprevir
HCV NS3/4A: hepatitis C virus nonstructural protein 3/4A protease
KDPG: 2-keto-3-deoxy-6-phosphogluconate
MCP: MS2 bacteriophage coat protein
MOI: multiplicity of infection
P2A/T2A: porcine teschovirus-1 2A and Thosea asigna virus 2A peptide
PH: pleckstrin homology
PI(4,5)P₂: phosphatidylinositol 4,5-bisphosphate
VPS4A: vacuolar protein sorting-associated protein 4A
VSV-G: vesicular stomatitis virus glycoprotein G
λN: lambda bacteriophage N peptide

## Acknowledgments

We thank Kanae Sasaki for helpful comments on the manuscript; Sar Lindner at the Harvard Medical School Electron Microscopy Facility for assistance with EV negative staining and transmission electron microscopy; and the DRSC/TRiP Functional Genomics Resources and DRSC-BTRR at Harvard Medical School for access to the SpectraMax Paradigm multi-mode microplate reader. This work was conducted with consulting service for statistical analyses from UM1TR004408 award through Harvard Catalyst | The Harvard Clinical and Translational Science Center (National Center for Advancing Translational Sciences, National Institutes of Health) and financial contributions from Harvard University and its affiliated academic healthcare centers.

