## Supplementary material for "Humanized Extracellular Vesicles for Efficient RNA Delivery": SI Appendix Materials and Methods

Plasmids, antibodies, and other reagents. EVPC, EVP, and EVC plasmids were constructed by Gibson assembly. The rat PLCδ1 PH domain, I3-01 nanocage, and HIV-1 Gag p6 elements were PCR-amplified from pCMV-EPN-24 (Addgene #79939). HCV NS3/4A (with its cleavage site) was amplified from pEF NS34a-msGFP-'Q-mito (Addgene #158764). Tandem BoxB arrays were amplified from prCAG-BoxB (Addgene #191122) and pCMV5-25BoxB (Addgene #60817). Tandem λN repeats, human CLYBL, human CEP55 EABR, and human epsin1 ENTH sequences were synthesized as oligonucleotides or double-stranded gene fragments. pCALSL-DsRed was generated by replacing the NeoR/KanR cassette in pCALSL-DsRed (Addgene #13769) with the triple SV40 poly (A) signal from Ai9 (Addgene #22799). The VSV-G (Indiana strain) expression plasmid was a kind gift from Dr. Christopher L. Parks. The pEGFP-VPS4-E228Q plasmid was obtained from Addgene (#80351). All new constructs were verified by whole-plasmid sequencing. For immunoblotting, the primary anti-Myc antibody was mouse monoclonal clone 4A6 (Sigma-Aldrich, 05-724); the secondary antibody was HRP-conjugated anti-mouse IgG (sheep) (Amersham/Cytiva, NA931). Grazoprevir (MK-5172; HY-15298) and α-amanitin (HY-19610) were obtained from MedChemExpress. BenzonaseÂ® Nuclease (E1014; Sigma-Aldrich) was used as indicated. Plasmid transfections were performed with Lipofectamine™ 2000 (11668019; Thermo Fisher Scientific) or linear polyethylenimine (PEI, 25 kDa) (23966; Polysciences) according to the manufacturers’ instructions.

Cell lines. 293T cells (ATCC® CRL-3216™) and BHK-21 cells (EH1011, Kerafast) were cultured in Dulbecco’s modified Eagle’s medium (DMEM, high glucose, pyruvate, 11995065, Thermo Fisher Scientific) supplemented with 10% heat-inactivated fetal bovine serum (FBS, 10437028, Thermo Fisher Scientific) and 1 U/ml penicillin-streptomycin (15140-163, Invitrogen) at 37 °C and 5% CO2.

EV production. 293T cells were split 1:3 into 24-well plates. After 24 h, cells were transfected with either EVPCs (200 ng/well) or EVP + EVC (200 ng each/well) together with VSV-G (100 ng/well) using Lipofectamine™ 2000 (2.5 μL/well). In each batch, one group was transfected with either EVPC-5 alone or with EVP+EVC constructs in the absence of VSV-G (Non-VSV-G control). Twenty-four hours post-transfection, the medium was replaced with 1 mL DMEM containing 2% exosome-depleted FBS (A2720803, Thermo Fisher Scientific). For constructs encoding HCV NS3/4A (wild type or S139A inactive variant), the production medium was supplemented with grazoprevir (1,250 nM) unless otherwise noted (Fig. 1G). Supernatants were harvested 48 h after the medium change for downstream analyses immediately or stored at -80 °C.

EV titration assay. BHK-21 cells were split 1:8 into 24-well plates. After 24 h, cells were transfected with pCALSL-DsRed (200 ng/well) using linear PEI at a DNA:PEI mass ratio of 1:6. Three hours post-transfection, the medium was replaced with 0.5 mL DMEM. EV-containing supernatants were pretreated with Benzonase® Nuclease (≥250 U/µL) at 0.1 µL per 100 µL supernatant (in the presence of 1 mM MgCl₂) and incubated at 37 °C for 30 min. EV-containing supernatants were serially diluted 10-fold across the wells. In the first well, 10 µL of undiluted supernatant was added. One transfected well without EV addition served as the negative control. After 4 days, random fields from the first well were imaged. Prior to counting, the excitation/laser power on the inverted fluorescence microscope was reduced until fewer than 10 DsRed⁺ cells, attributable to leaky expression from pCALSL-DsRed, were visible in the negative-control well. Counts were then performed on wells containing fewer than 2,000 DsRed⁺ cells. Functional titer (DU/mL) was calculated as: DU/mL = 𝑁× multiplier, where N is the number of DsRed⁺ cells in that well, the multiplier is 100 for the first well, 1000 for the second well, and so on.

Western blot. EV-containing supernatants (5.5 mL) from WT EVPC-5 and EVPC-5 (S139A) were diluted to 33.5 mL in DMEM and concentrated by sucrose-cushion ultracentrifugation over a 5 mL, 20% sucrose cushion using a swinging-bucket SW 32 Ti rotor at 21,000 rpm for 1.5 h (single batch) or 2 h (two batches). The EV pellets were collected in the bottom of the tubes and resuspended in 50 µL of DMEM, briefly vortexed and centrifuged, then incubated at 37 °C for 30 min to permit NS3/4A cleavage. Concentrated EVs (20 µL) were mixed with 20 µL Laemmli buffer (2×) and heated at 95 °C for 5 min. Samples were resolved on 4–12% Bolt™ Bis-Tris Plus Mini Protein Gels with MOPS running buffer. Anti-Myc (1:2000) was used to detect the I3-01–Myc tag; blots were developed with HRP-conjugated anti-mouse IgG (1:5000) and Pierce™ ECL Western Blotting Substrate (32109, Thermo Fisher Scientific).

RT–qPCR quantification of exported mRNA. Quantification was adapted from ([Addgene: AAV Titration by qPCR](https://www.addgene.org/protocols/aav-titration-qpcr-using-sybr-green-technology/)) originally developed for AAV titers. The mRNA copy number per mL in supernatants was determined by RT–qPCR using a standard curve generated from linear dsDNA corresponding to Cre–P2A–EGFP, excised from EVC-2 by NheI/BsrGI digestion. The dsDNA concentration was measured with the AccuGreen™ Broad Range dsDNA Quantitation Kit (Biotium, 31069). A standard curve (four points, 1:10 serial dilutions) was run on every plate. Total RNA from 100 µL of supernatant was isolated with the Direct-zol RNA Miniprep Kit (Zymo Research, R2052) following the manufacturer’s instructions and eluted in 25 µL H₂O. 4 µL of RNA were used to synthesize first-strand cDNA with SuperScript III Reverse Transcriptase (Thermo Fisher Scientific, 18080085) in a 20 µL reaction, using a gene-specific primer targeting Cre (5′-GAGTCCAGGTTTCTGATGTAGTTC-3′). qPCR was performed with PowerUp™ SYBR™ Green Master Mix (Thermo Fisher Scientific, A25780) on a CFX96 real-time PCR instrument (BioRad). Primers were modified from Horns et al. (11): F 5′-CCAACAACTACCTGTTCTGC-3′ and R 5′-GCCTCAAAGATCCCTTCCAG-3′. cDNA was diluted 20-fold prior to qPCR. Each sample was run in triplicate; data were accepted when at least two replicates showed ΔCt < 0.5. Otherwise, the qPCR was repeated. Copy numbers per reaction were converted to copies/mL of supernatant using the standard-curve fit and the recorded dilution factors.

NanoLuc assays following EV delivery. EV-containing supernatants (100 µL) were pretreated with Benzonase® Nuclease and incubated at 37 °C for 30 min, as in the titration assay. 293T cells at ~100% confluency in 6-well plates served as recipients. Culture medium was replaced with 5 mL DMEM, and 100 µL of the treated supernatant was added (permitting dilution of grazoprevir, when present). After 3 h, the medium was replaced with 2 mL DMEM + 10% FBS containing 1 µg/mL α-amanitin. Because residual plasmid DNA can confound NanoLuc readouts intended to measure translation from EV-delivered mRNA, Benzonase pretreatment and RNA polymerase II inhibition were included. After 22 h, cells were harvested in 500 µL Passive Lysis Buffer (E1941, Promega), and 50 µL aliquots were assayed in triplicate using the Nano-Glo Luciferase Assay (N1110, Promega) according to the manufacturer’s instructions on a SpectraMax Paradigm multi-mode microplate reader (Molecular Devices). For each batch, NanoLuc activity in every group was normalized to the mean of three biological replicates from the EVP-4 + EVC-6 group.

NanoLuc assays following plasmid transfection. 293T cells were transfected in 24-well plates using the same protocol as for EV production, except VSV-G was omitted. Each combination was performed in duplicate. After 22 h, cells were lysed in 50 µL Passive Lysis Buffer (E1941, Promega), and lysates from duplicates were combined. Pooled samples were diluted 80- or 100-fold in Passive Lysis Buffer, and 50 µL aliquots were assayed in triplicate using the Nano-Glo Luciferase Assay (Promega), as described for the NanoLuc assays following EV delivery. For each batch, NanoLuc activity in all groups was normalized to the EVP-4 + EVC-6 group within the same batch.

Electron microscopy. EVs were concentrated by sucrose-cushion ultracentrifugation over a 5 mL, 20% sucrose cushion using a swinging-bucket rotor, SW 32 Ti rotor, at 21,000 rpm for 1.5 h. The pellets were collected in the bottom of the tubes and resuspended in 100 µL of DMEM. A 5 µL aliquot of the sample (either neat or 1:1 diluted with ddH₂O) was adsorbed for 1 minute to a carbon coated grid that had been made hydrophilic by a 20 second exposure to a glow discharge (25mA). Excess liquid was removed with a filterpaper (Whatman #1), the grid was then floated briefly on a drop of water (to wash away phosphate or salt), blotted again on a filer paper and then stained with 1% Uranyl Acetate (EMS catalog # 22400) for 20-30 seconds. After removing the excess stain with a filter paper the grids were examined in a JEOL JEM-120i Transmission electron microscope and images were recorded with an NS15 camera.

Statistical analysis. Data are presented as mean ± SEM. Statistical analyses were performed using GraphPad Prism. Sample sizes are shown in the figures, and the statistical tests applied to each experiment are described in the corresponding figure legends. Normality was assessed using the Shapiro–Wilk test, and homogeneity of variances was evaluated using the Brown–Forsythe test. For two-group comparisons, if data were normally distributed, unpaired t-tests (with Welch’s correction when variances were unequal) or paired t-tests (for matched data) were applied. When the assumption of normality was not met, nonparametric alternatives were used: the Mann–Whitney test for unpaired data or the Wilcoxon signed-rank test for paired data. For comparisons involving more than two groups without repeated measures, when the assumption of normality was not met, the nonparametric Kruskal–Wallis test with Dunn’s multiple comparisons was applied. When the data were normally distributed but variances were unequal, Welch’s ANOVA was applied, followed by Dunnett’s T3 multiple comparisons test. When both assumptions were satisfied, one-way ANOVA followed by Dunnett’s or Tukey’s multiple comparisons test was used. For comparisons involving more than two groups with repeated measures, when the assumption of normality was satisfied, repeated-measures one-way ANOVA with the Geisser–Greenhouse correction was performed, followed by Dunnett’s multiple comparisons test. When the assumption of normality was not met, the Friedman test for repeated-measures designs was performed, followed by Dunn’s multiple comparisons test. For comparisons involving a single group against a reference value, one-sample t-tests were performed, as all groups followed the normal distribution. Significance thresholds: ns (not significant) *p* ≥ 0.05, ✱ *p* < 0.05, ✱✱ *p* < 0.01, ✱✱✱ *p* < 0.001.

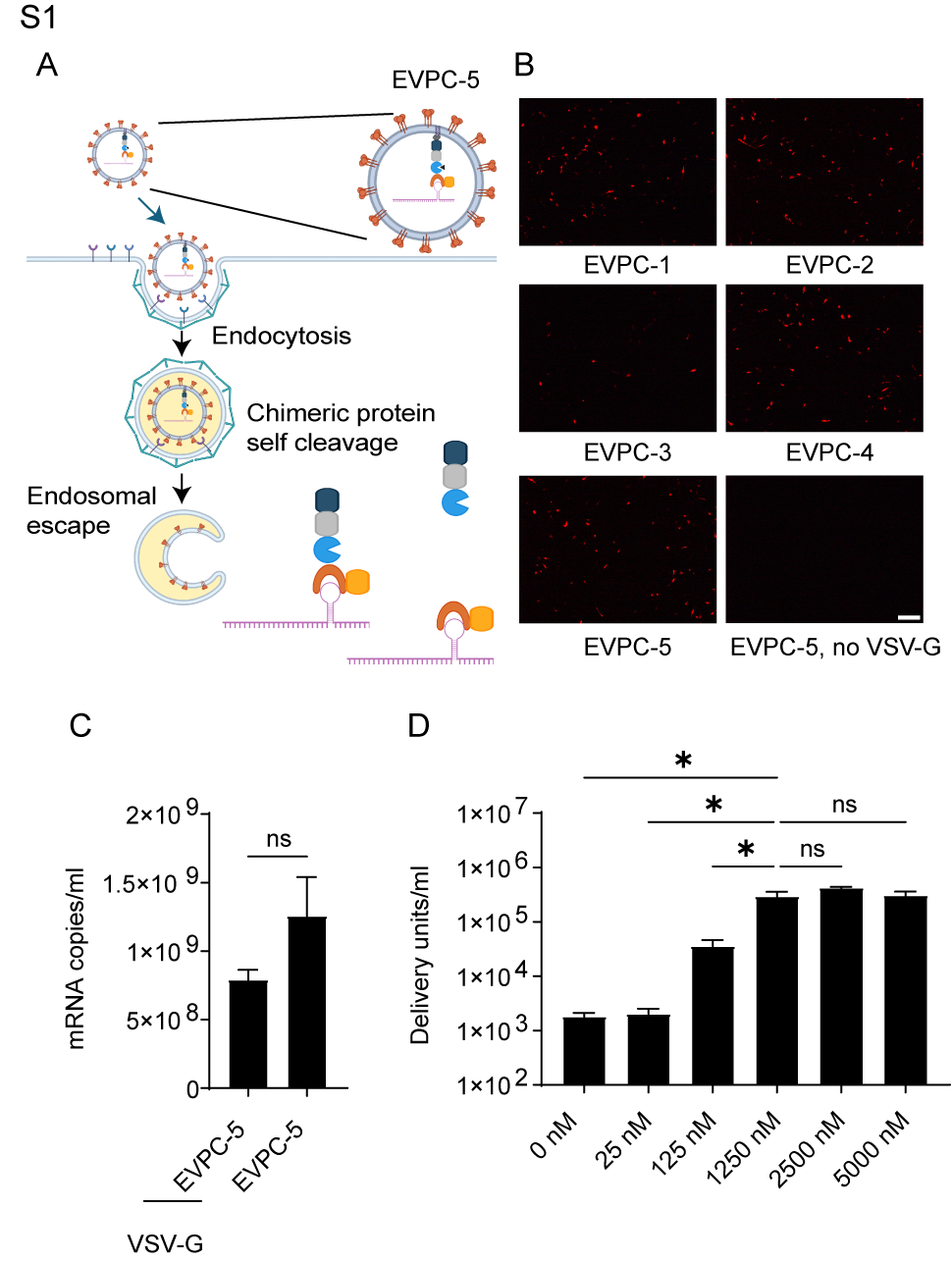

Fig. S1 related to Fig. 1.

(A) Protease-modulated EV mRNA release in EVPC-5. During EV production, GZR inhibits NS3/4A protease, enabling efficient mRNA loading and EV biogenesis. Upon VSV-G mediated endocytosis and endosomal escape in recipient cells, dilution of GZR allows NS3/4A to cleave at its C-terminal site, releasing the λN-tethered mRNA from the PH–I3-01 nanocage to promote translation.

(B) Representative fluorescent images of DsRed⁺ BHK-21 recipient cells from the first dilution well (1:50) of the EV titration assay related to Figure 1D. Scale bar, 100 μm.

(C) Test of VSV-G as a requirement for functional mRNA delivery. EVPC-5 was transfected without VSV-G despite similar levels of exported Cre mRNA in producer supernatants by RT-qPCR (mean ± SEM; n = 5). *p*-value was calculated using a paired t-test.

(D) GZR dose–EV titer curve. Recipient cells were treated with different concentrations of GZR to determine the concentration needed for effective release from protease inhibition (n = 5). *p*-values were calculated using Dunnett’s multiple comparison test following repeated-measures one-way ANOVA with the Geisser–Greenhouse correction.

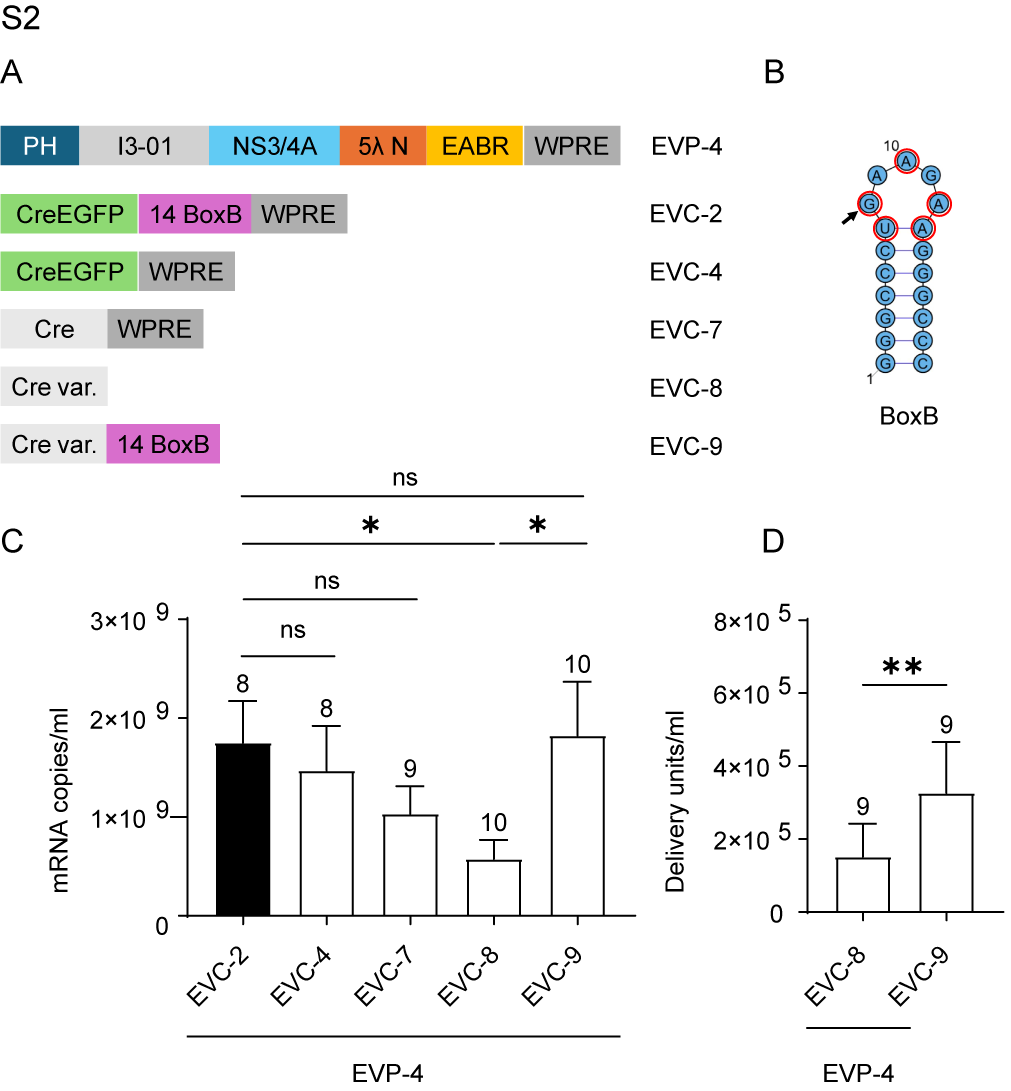

Fig. S2. Effect of BoxB mimic sequence in mRNA loading and functional delivery in combination with EVP-4.

(A) Design schematic.

(B) Schematic of the 19-nt BoxB hairpin (7-bp stem, 5-nt loop) with the loop consensus UGNANAA; key loop nucleotides are circled in red. The G8 nucleotide mutated in EVC-8 is indicated by an arrow.

(C) mRNA export levels in five groups with stepwise removal of BoxB/BoxB-mimic motifs and reintroduction of 14×BoxB. *p*-values were calculated using Kruskal–Wallis test with Dunn’s multiple comparisons vs EVC-2 group. Comparison between EVC-8 and EVC-9 was performed using the Wilcoxon test.

(D) EV functional titers in two groups showed in (C). *p*-value was calculated using the Wilcoxon test.

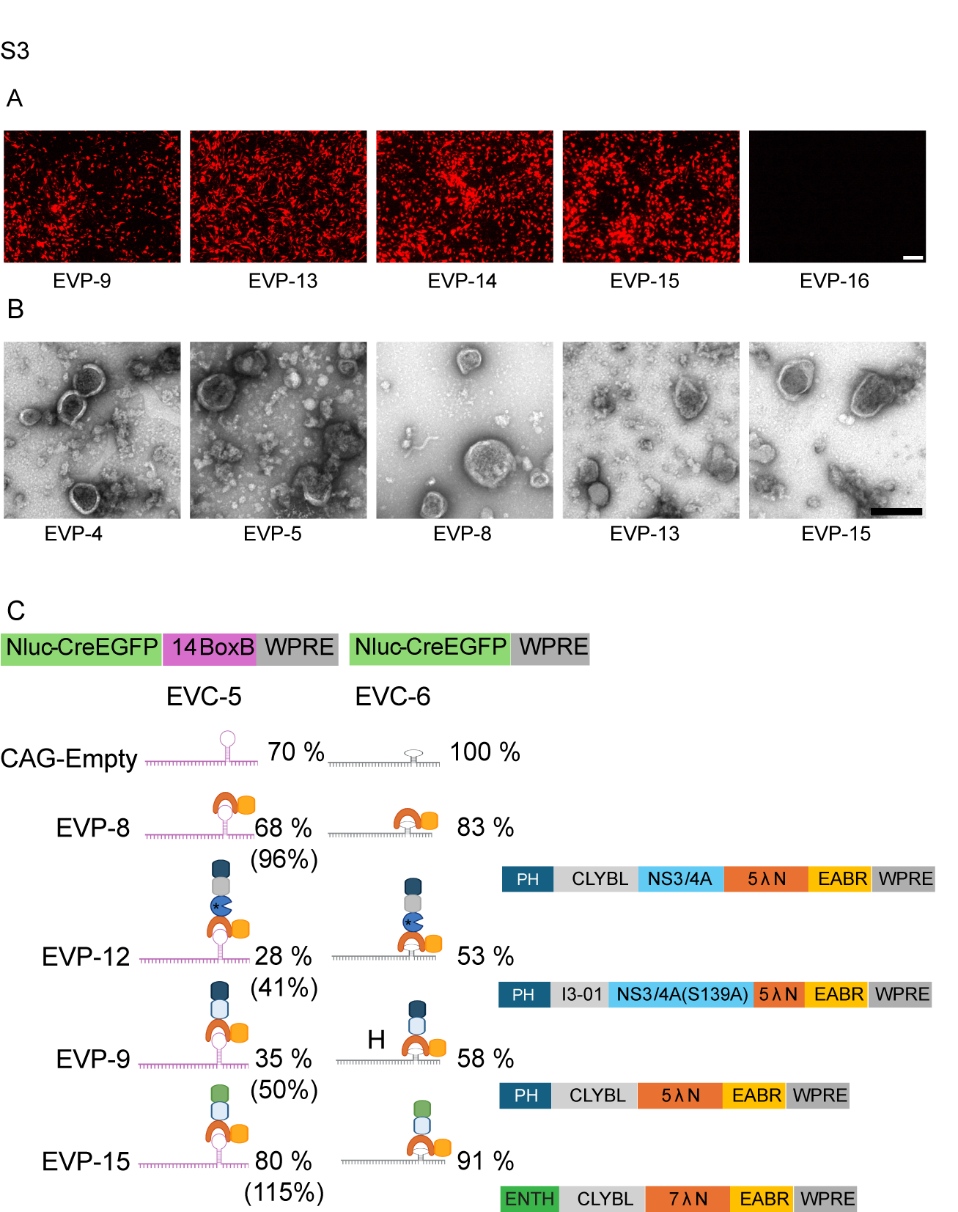
.

Fig. S3 related to Fig. 6 and Fig. 7.

(A) Representative fluorescent images of DsRed⁺ BHK-21 recipient cells from the first dilution well of the EV titration assay for the six groups in Figure 7A. Scale bar, 100 μm.

(B) Negative-stain EM of EVs from EVPs with EVC-6. Scale bar, 200 nm.

(C) Diagram of combined datasets (part of Fig. 6H and Fig. 7E). NanoLuc from EVP/EVC co-transfections of 293T (EVC-5 or EVC-6) was (i) normalized to EVP-4 + EVC-6, then (ii) rescaled so that the average CAG-empty + EVC-6 = 100%; all values are shown as % of CAG-empty + EVC-6. For EVC-5 combinations, a second value—% of CAG-empty + EVC-5—is provided in parentheses.
