## Supplementary figures and images for "Humanized Extracellular Vesicles for Efficient RNA Delivery"

### SI Appendix Fig. 1

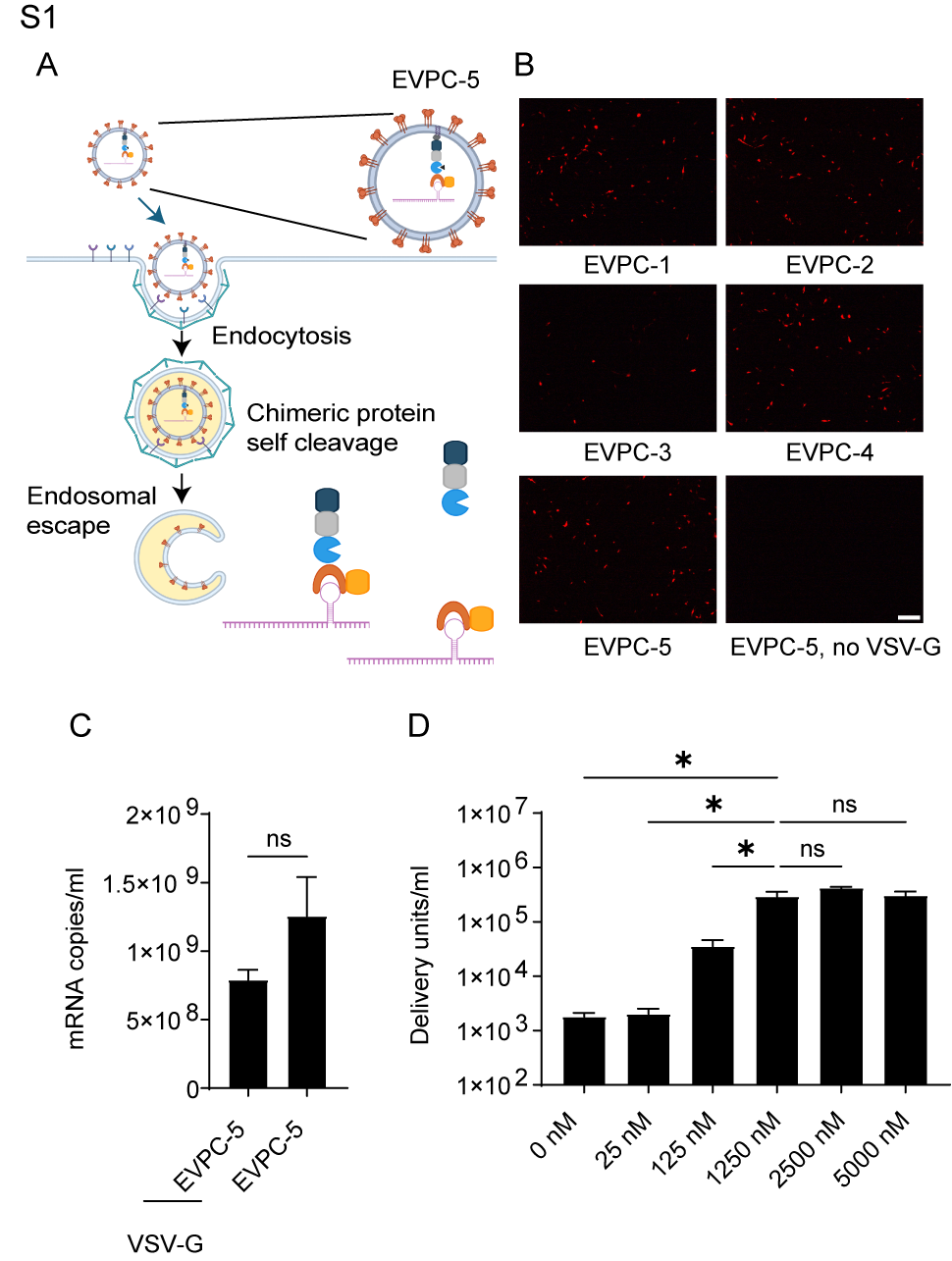

### SI Appendix Fig. 2

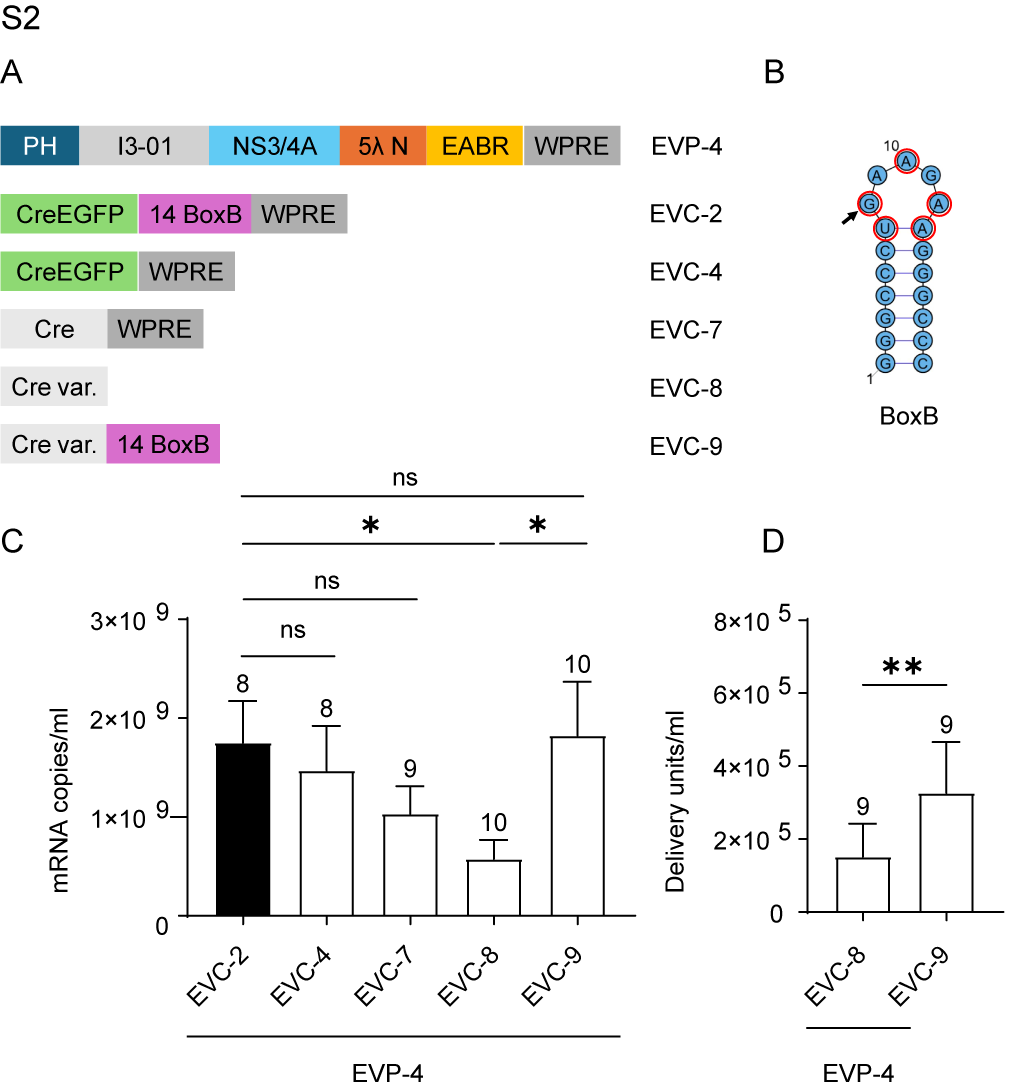

### SI Appendix Fig. 3

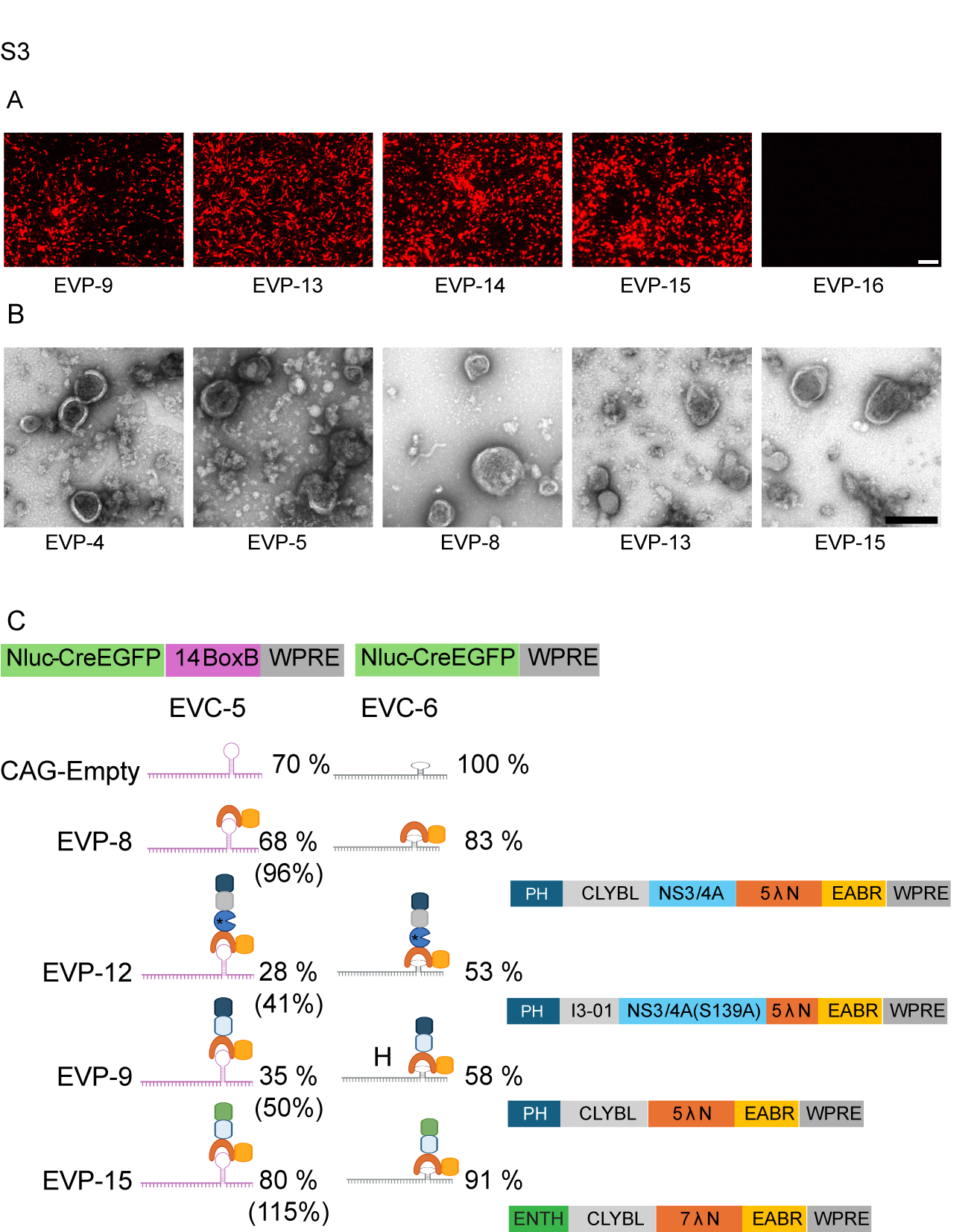
